# Bromodomains regulate dynamic targeting of the PBAF chromatin remodeling complex to chromatin hubs

**DOI:** 10.1101/111674

**Authors:** C.A. Kenworthy, N. Haque, S.H. Liou, P. Chandris, V. Wong, P. Dziuba, L.D. Lavis, W.L. Liu, R.H. Singer, R.A. Coleman

**Affiliations:** Albert Einstein College of Medicine; Gruss-Lipper Biophotonics Center, Department of Anatomy and Structural Biology, Albert Einstein College of Medicine, Bronx, New York 10461, USA; Section on High Resolution Optical Imaging, National Institute on Biomedical Imaging and Bioengineering, National Institutes of Health, Bethesda, Maryland, 20892, USA; HHMI Janelia Research Campus; Albert Einstein College of Medicine, HHMI Janelia Research Campus

**Keywords:** PBAF, chromatin remodeling, Histone acetylation, Histone H3.3, Single Molecule Tracking, bromodomain, nuclear hubs

## Abstract

Transcriptional bursting involves genes rapidly switching between active and inactive states. Chromatin remodelers actively target arrays of acetylated nucleosomes at select enhancers and promoters to facilitate or shut down the repeated recruitment of RNA Pol II during transcriptional bursting. It is unknown how acetylated chromatin is dynamically targeted and regulated by chromatin remodelers such as PBAF. Thus, we sought to understand how PBAF targets acetylated chromatin using live-cell single molecule fluorescence microscopy. Our work reveals chromatin hubs throughout the nucleus where PBAF rapidly cycles on and off the genome. Deletion of PBAF’s bromodomains impairs targeting, stable engagement and persistent binding on chromatin in hubs. Interestingly, PBAF has a higher probability to stably engage chromatin inside hubs indicating that hubs contain a unique nucleosomal scaffold compared to global chromatin. Dual color imaging of PBAF in hubs near H3.3 or HP1α reveals that PBAF targets both euchromatic and heterochromatic regions with distinct genome binding kinetics that mimic chromatin stability. Removal of PBAF’s bromodomains stabilizes H3.3 and HP1α binding within chromatin indicating that bromodomains may play a direct role in remodeling of the nucleosome. Our data, suggests that PBAF differentially and dynamically engages a variety of chromatin structures involved in both activation and repression of transcription via bromodomains. Furthermore, PBAF’s binding stability on chromatin may reflect the chromatin remodeling potential of different bound chromatin states.

**Statement of Significance:** Transcriptional bursting involves a gene rapidly switching between transcriptionally active and inactive states. To regulate transcriptional bursting, chromatin must interchange between euchromatin and heterochromatin to permit or restrict access of transcription factors including RNA Polymerase II to enhancer and gene promoters. However, little is known regarding how chromatin remodelers dynamically read a rapidly changing 4D epigenome. We used live-cell single molecule imaging to characterize the spatiotemporal chromatin binding dynamics of PBAF, a chromatin remodeler that accesses both euchromatin and heterochromatin to regulate transcription. PBAF cycles on and off chromatin hubs in select nuclear regions where it distinctly engages euchromatin and heterochromatin via bromodomains in its BAF180 subunit. Our study provides the framework to understand how the 4D epigenome is regulated.

## Introduction

Transcriptional bursting is defined by brief periods of time where a gene promoter is in a highly permissible state that allows transcription factor recruitment and rapid loading of RNA Polymerase II (RNA Pol II) every 8-20 seconds [1]. After a period of minutes, the gene is shut down and transcriptionally silent for a period of minutes to hours [2]. There is a tight linkage between transcriptional activity and chromatin state changes such as the interconversion between euchromatin and heterochromatin. Therefore, it is likely that chromatin remodelers, histone variants and factors that actively stabilize different chromatin states also bind and unbind enhancers and promoters in a cyclical manner.

Recent live-cell imaging studies indicate that transcription factors (e.g. RNA Pol II, Mediator and Sox2) dynamically bind chromatin as clusters to form hubs of activity that regulate local gene expression [3–8]. However, such dynamic activity of chromatin modifiers within distinct chromatin hubs is currently poorly characterized due to a number of technical limitations. In particular, researchers lack efficient methods to identify and quantitatively characterize chromatin in active/inactive hubs in live cells.

PBAF is ATP-dependent chromatin remodeling complex that both evicts or repositions nucleosomes to regulate transcription of stress response genes via bromodomain-dependent targeting of acetylated chromatin [9–11]. Numerous in vitro studies have found that removal or mutational inactivation of bromodomains in the BAF180 subunit reduces PBAF’s binding to chromatin [12, 13]. However, none of these studies determined if bromodomains regulate both the association and dissociation of PBAF with acetylated chromatin in vivo. In addition, PBAF targets heterochromatin to actively repress transcription likely via repositioning or stabilizing a nucleosome at promoters and enhancers through a poorly characterized mechanism [14–17]. Therefore understanding how PBAF dynamically recognizes highly localized hubs of different chromatin states may lead to a greater understanding of the spatial and dynamic regulation of chromatin topology and gene regulation *in vivo*.

To spatially distinguish and characterize different types of chromatin hubs *in vivo*, we used live-cell <u>S</u>ingle <u>M</u>olecule <u>T</u>racking (SMT) to dynamically map chromatin binding of PBAF alongside prototypical markers of euchromatin (H3.3) and heterochromatin (HP1α). Our dynamic imaging studies reveal small hubs where PBAF cycles on and off chromatin. To map out different chromatin states, we investigated PBAF’s engagement and stability on chromatin when encountering H3.3 and HP1α marked hubs. More importantly, we have assessed the role of PBAF’s bromodomains in hub targeting, cycling on chromatin and select engagement with different types of chromatin hubs. Overall, our studies provide new insights into dynamic chromatin targeting of PBAF via bromodomains.

## Materials and Methods

### Plasmid constructions, generation of cell lines and live-cell fluorescent labeling of proteins

Details of plasmid construction, generation of cell lines and fluorescent labeling of proteins are described in the Supplemental Methods section.

### Live-cell single molecule imaging of Halo-BAF180 WT/ΔBD, H3.3-SNAP or SNAP-HP1α

All imaging sessions were carried out at room temperature in L-15 media (Gibco) to support cell growth in conditions lacking CO_2_. Experiments were performed at room temperature to minimize microscope drift and cell movement. Imaging experiments performed at 37°C showed equivalent residence times and distributions of Halo-BAF180 compared to imaging performed at room temperature. Cells were continuously illuminated using a 532 nm (13 W/cm^2^, Coherent) or 640 nm (9.5 W/cm^2^, Coherent) laser for JF549-HTL and SNAP-Cell 647-SiR imaging respectively. Time-lapse two-dimensional (2D) images of single molecules were acquired with a customized inverted Nikon Eclipse Ti microscope with a 100X oil-immersion objective lens (Nikon, 1.49 NA) and further magnified 1.9X post-objective. BAF180 images were acquired at 2 Hz for ~18 minutes using an EMCCD (iXon, Andor) with a 512 x 512 pixel field of view (final pixel size of 84 nm). SNAP imaging proceeded at 2 Hz for ~4.5 minutes in cells that expressed either SNAP-Cell 647-SiR labeled H3.3-SNAP or SNAP-HP1α.

### Image processing and single molecule tracking

Movies of acquired images were processed to subtract background in ImageJ using a rolling ball radius of 50 pixels. Background subtracted movies were subjected to <u>M</u>ulti-<u>T</u>arget <u>T</u>racking (MTT) to resolve the trajectories of individual molecules [39] using a GUI based implementation, SLIMfast [40]. Localization of individual molecules was achieved by fitting Point Spread Functions (PSFs) of discrete single spots with a 2D gaussian function. Tracking of single molecule chromatin binding events was performed by connecting BAF180 localizations between consecutive frames. Tracking was based upon a maximum expected diffusion constant of 0.05 μm^2^/second and allows for 1.5 second gaps in trajectories due to blinking or missed localizations. Track positions from individual trajectories were averaged to generate a 2D projection map of BAF180 binding events over 18 minutes of imaging.

Nuclear BAF180 tracks were identified based on the boundaries from 2D projection maps of 180 binding events. BAF180 tracks that fell outside of the nucleus were excluded. Photobleach rates were then determined for each background-subtracted movie based upon exponential decay of the global fluorescence of chromatin bound Halo-BAF180 WT/ΔBD, H3.3-SNAP, SNAP-HP1α.

### Analysis of PBAF chromatin binding residence times

Chromatin binding residence time was determined by plotting a survival curve (1-<u>C</u>umulative <u>D</u>ensity <u>F</u>unction, 1-CDF) of the track-lengths of chromatin bound Halo-BAF180 in each cell. Single and double-exponential models were then fitted to these 1-CDF plots to determine residence times. One-way ANOVA followed by Tukey’s post-hoc t-tests were performed to determine pairwise significance of global residence times and percentages of stable PBAF binding events.

### Analysis of PBAF clustering in hubs

2D projection maps of BAF180 binding events lasting at least one second (Figures 2E and 2F) or eight seconds (Figure 2C) were expanded 10 fold in the X and Y directions yielding a final pixel size of 8.4 nm. Areas of high PBAF binding densities were determined by counting the number of binding events within an octagon window (diameter 168 nm) as it was raster scanned across the nucleus of the expanded 2D projection map. Contiguous octagon widows centered on an individual pixel containing at least 3 PBAF binding events were defined and labeled as hubs. The total number of hubs per cell were then normalized to the total PBAF binding events per cell and multiplied to produce the number of PBAF hubs formed per 10,000 PBAF binding events (i.e. Number of PBAF hubs) for each cell. The median number of PBAF binding events inside hubs in cells was also normalized to the total PBAF binding events per cell and multiplied to produce the number of PBAF binding events in hubs per 10,000 PBAF binding events over 18 minutes of imaging. Overall significance was determined with a two-sample Kruskal-Wallis test to determine pair-wise significance.

**Figure 1:**
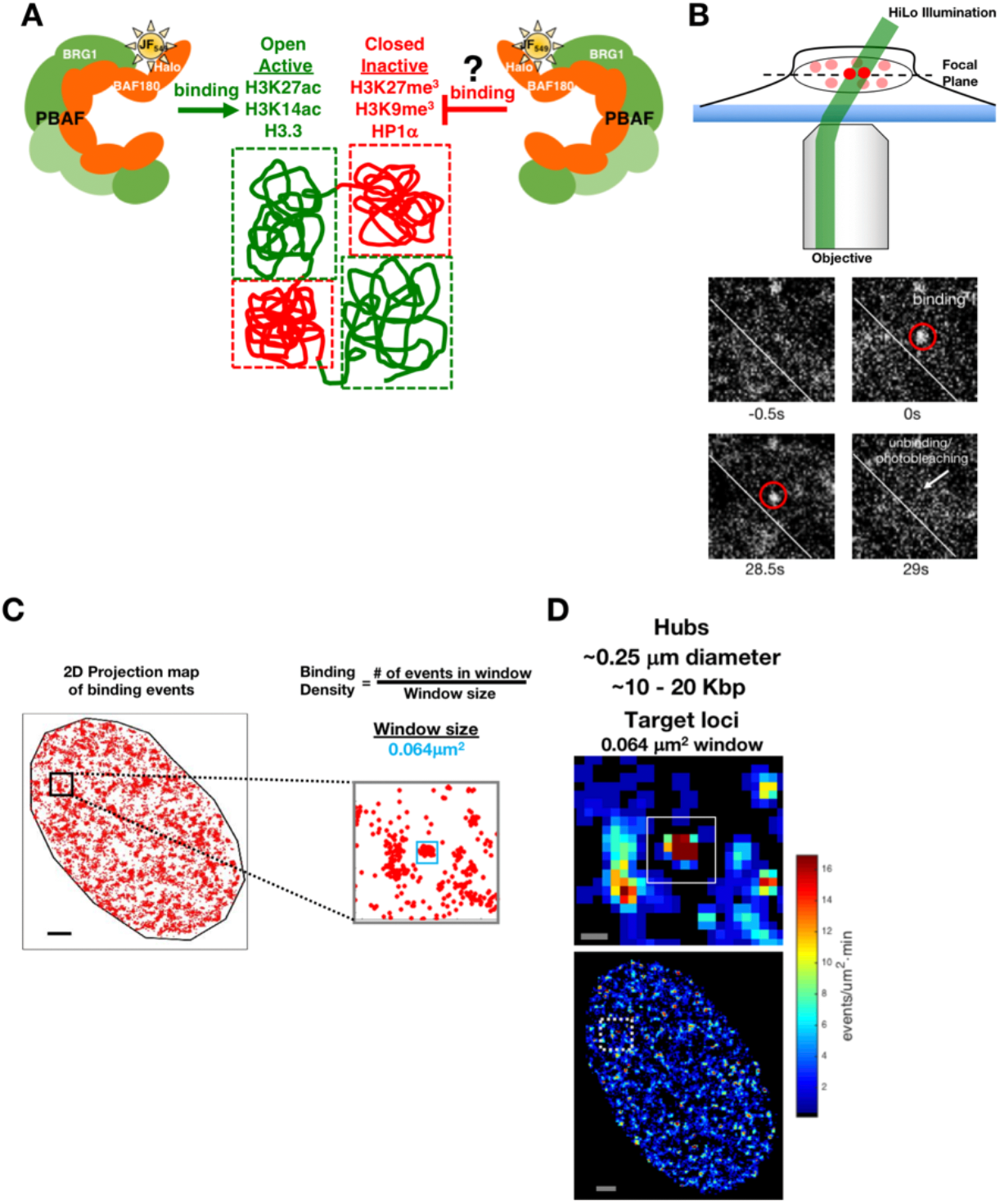
Spatial analysis of PBAF chromatin binding events using SMT to define binding hubs. *(A)* Transcriptionally active genomic regions (green) contain open chromatin structures associated with acetylated histone marks (e.g. H3K27ac & H3K14ac) and the H3.3 histone variant. Transcriptionally inactive genomic regions (red) contain closed chromatin structures associated with select methylated histone marks (e.g. H3K27me^3^ & H3K9me^3^) and heterochromatic protein HP1α. PBAF contains multiple bromodomains within the BAF180 subunit known to bind acetylated histone marks associated with open chromatin regions. *(B)* Motion Blur HiLo microscopy (top panel) of a single U2OS cell stably expressing Halo-BAF180 WT. PBAF containing Halo-BAF180 WT molecules that rapidly diffuse in the nucleoplasm are blurred, while chromatin bound PBAF appears as single bright spots (highlighted by red circles, lower panels). Disappearance of a spot (white arrow) is due to unbinding or dissociation of PBAF from the chromatin. *(C)* Strategy for measuring the non-homogeneous localization of PBAF chromatin binding events in a nucleus. A 2D projection map of PBAF binding events (red dots) in the nucleus over 18 minutes of imaging is shown. A grey box (left panel, zoomed view in the right panel) outlines a representative window of PBAF binding events in a subnuclear region. PBAF binding density is thereby determined by counting the number of binding events located within a given size window (0.064 μm^2^, blue box). Scale bar, 2 μm. *(D)* PBAF binding event frequency heat maps were obtained using the 2D projection map in *(C).* Regions of high (red) and low (blue) PBAF binding frequency is presented for the cell shown in *(C).* The top panel is a zoomed in view of the dashed box in lower panel. PBAF hubs (top left panel, white box) were identified as clusters of frequent PBAF chromatin binding to target loci. Scale bar, 0.25 μm (top panel) and 2 μm (bottom panel), respectively.

**Figure 2:**
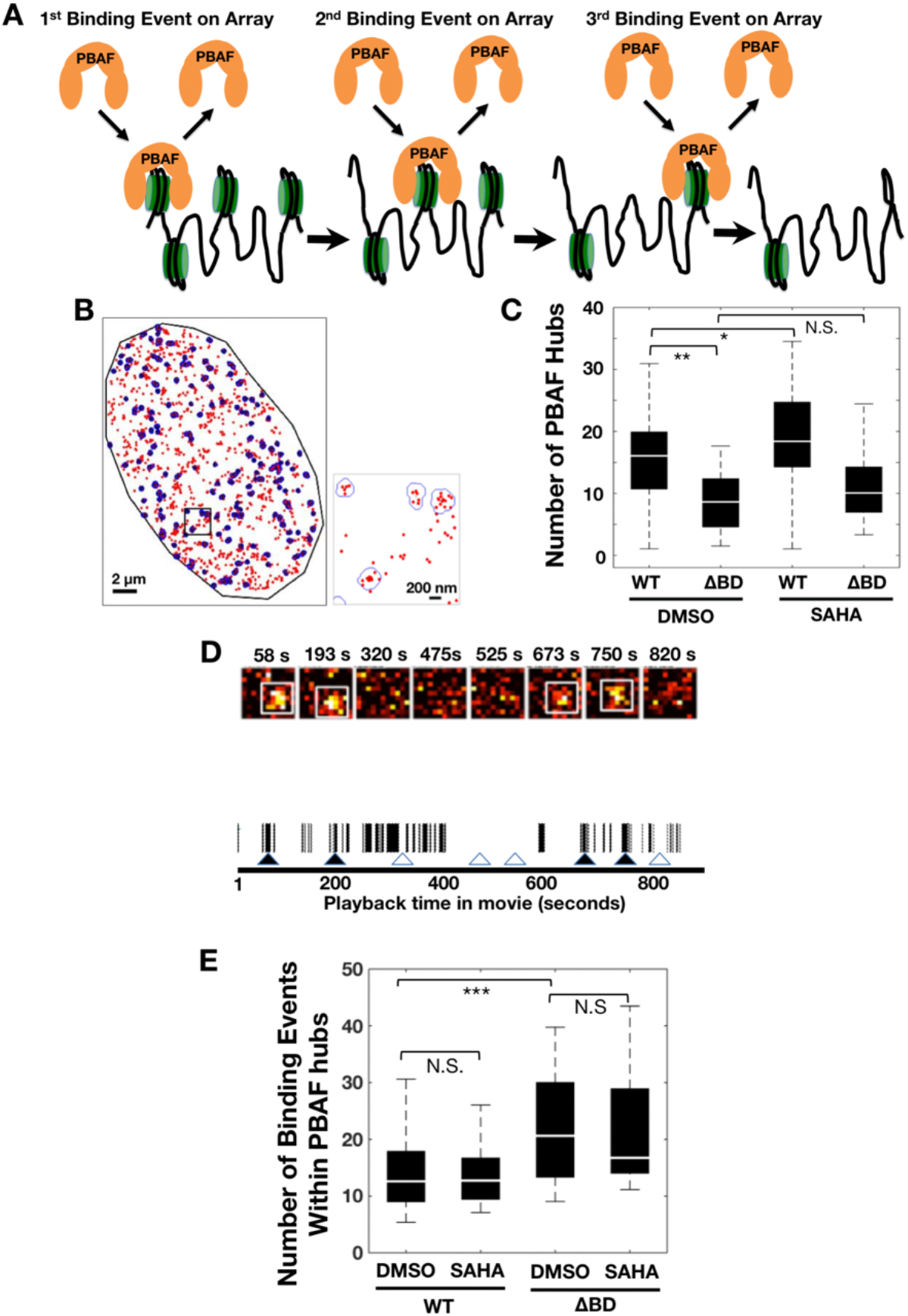
Spatial and cycling analysis of PBAF binding hubs *(A)* A schematic representation where PBAF cycles on and off an array of nucleosomes. *(B)* Clustering analysis algorithms indicate PBAF binding events (red dots) within hubs (left panel, blue outlines). The right panel displays an expanded inset of a boxed region in the left panel. *(C)* The number of hubs formed per cell, which express either wild-type PBAF WT or the mutant PBAF ΔBD. N.S. = Not significant, p-value * = < 0.05 and ** = < 0.01. (N=39 WT DMSO cells, 17 ΔBD DMSO cells, 37 WT SAHA cells, and 21 ΔBD SAHA cells). *(D)* Temporal occupancy of PBAF at a representative chromatin binding hub. Snapshots of PBAF binding foci at different timepoints in movie (seconds) and the appearance of PBAF binding highlighted by white boxes (top panel). The width of the black bars represents the duration of individual PBAF/chromatin binding events in a hub (bottom panel). White gaps represent latent time periods when the region containing a hub is not occupied by PBAF. Timepoints (▲) from the top panel where PBAF binds in the hub. Timepoints (△) where PBAF does not bind the hub. *(E)* Median number of PBAF binding events per hub per cell (N=39 WT DMSO cells, 17 ΔBD DMSO cells, 37 WT SAHA cells, and 21 ΔBD SAHA cells). N.S. = Not significant and p-value * = < 0.05. p-value ** = < 0.01 and *** = < 0.001. For data in panels *(C)*, *(E)* and *(F)*, the white bar in the solid black box is the median, while the lower and upper boundaries of the black box correspond to the 25th and 75th percentiles of the data respectively. Outliers typically represented less than 10% of the dataset and therefore were omitted for clarity.

### Characterization of PBAF localization and binding dynamics in proximity to H3.3 or HP1α hubs

High binding density H3.3 or HP1α hubs were mapped using MTT and raster scanning as described above. Track lengths for PBAF molecules within 500nm of H3.3 or HP1α hubs were aggregated from a number of cells and plotted as a 1-CDF survival curve and fit to single- and double-exponential decay functions. The proximity distance of 500nm was chosen to reflect the size of the hubs (~200-400nm diameter) and the close association of PBAF with H3.3 or HP1α as seen in our fixed cell STORM microscopy experiments. Residence times of the specific binding population for PBAF in co-localized hubs were plotted as <u>P</u>robability <u>D</u>ensity <u>F</u>unctions (PDF). Statistical differences between genotype or treatment conditions were then assessed using a two-sample Kolmogorov-Smirnov test.

## Results

### PBAF targets chromatin in distinct nuclear hubs and compartments

A number of prior studies found that transcription factors, including EWS/FLI, Sox 2 and RNA Pol II, formed highly localized clusters or hubs of binding in enhancers or promoters within nuclear domains of ~200-300nm in diameter [4, 5, 7, 18]. However the molecular origins of dynamic transcription factor binding on chromatin hubs is poorly understood. Therefore, we chose to study how the rapid cycling of factors on and off of chromatin is influenced by histone post-translational modifications (PTMs) and chromatin subtypes (e.g. euchromatin versus heterochromatin).

Enhancers and promoters of transcriptionally active genes are enriched in acetylated chromatin. Accordingly, we hypothesized that chromatin remodelers such as PBAF, which targets a variety of acetylated residues via its 8-12 bromodomains [14, 19], would also form dynamic binding hubs in the nucleus (Figure 1A). To characterize the dynamic binding of PBAF to chromatin *in vivo*, we chose to fluorescently tag the BAF180 subunit (i.e. Halo-BAF180 WT, Figure S1), since it harbors six bromodomains critical for interaction with acetyl-lysine residues on histones [12, 13]. We confirmed that these Halo-tagged BAF180 proteins were incorporated into the large multi-subunit PBAF complex via co-immunoprecipitation studies and live-cell fast diffusion measurements (Supplemental Figures S1 and S2).

Motion Blur HILO microscopy combined with live-cell <u>S</u>ingle <u>M</u>olecule <u>T</u>racking (SMT) [20] was used to dynamically detect PBAF molecules bound to chromatin (Figure 1B and Supplemental Movie S1). At long camera exposure rates (~500 ms), fast-diffusing nuclear PBAF complexes are blurred out and cannot be localized. Single PBAF molecules, stably bound to chromatin, appear as distinct <u>P</u>oint <u>S</u>pread <u>F</u>unctions (PSFs) that are spatially and temporally resolved (Figure 1B). PSFs, representing PBAF’s binding and unbinding on chromatin, appear and disappear stochastically throughout the time course of imaging (Figure 1B and Supplemental Movie S1). Two dimensional (2D) projection maps showed select nuclear regions that contained high densities of PBAF chromatin binding events. (Figure 1C and Supplemental Figure S3C). Pair correlation function analysis, which was previously used for determination of transcription factor clustering [5], found that PBAF formed binding hubs of ~ 200-400nm in diameter (Supplemental Figure S3D).

To better identify and quantify PBAF’s dynamic binding in chromatin hubs, we developed an approach to spatially define the frequency of PBAF’s chromatin binding within nuclear subregions. These dynamic binding frequency heat maps were generated by raster scanning across the nucleus and counting the number of PBAF-chromatin binding events in a window of approximately 252nm diameter based on our pair correlation function analysis (Figures 1C and 1D). Spatially isolated regions spanning ~200-400nm diameter representing high frequency PBAF binding to chromatin were scattered throughout the nucleus (Figure 1D). No hubs were detected in simulations with random localizations of an equivalent number of binding events throughout the nucleus for every cell included in our analysis (see Supplemental Figure S4A for a representative cell). Based upon PBAF’s known role in remodeling chromatin in genomic elements associated with transcriptional regulation, we hypothesized that PBAF’s binding in equivalently sized hubs as Sox2 and RNA Pol II was due, in part, to dynamic interactions with chromatin in enhancers and promoters.

### PBAF targeting to chromatin hubs is regulated by BAF180 bromodomains

Histone PTMs associated with transcriptional regulation (e.g. acetylation) are ideal targets to better understand the molecular origins of PBAF cycling on chromatin hubs. Therefore, we chose to see how disruption of PBAF’s interaction with histone PTMs affected targeting to chromatin hubs. Arrays of acetylated nucleosomes can be repeatedly targeted by bromodomain containing chromatin remodeling complexes such as PBAF (Figure 2A) [21]. BAF180’s six bromodomains allow PBAF to recognize a large variety of acetyl-lysine residues in chromatin [19, 22–24]. Therefore, we compared the high frequency binding of wild-type PBAF (WT) and a mutant PBAF lacking the six BAF180 bromodomains (i.e. ΔBD) (Figure 2B and Supplemental Figure 1A). Non-specific probing of PBAF on chromatin could generate background noise that impairs hub detection. Thus, we limited our analysis to PBAF chromatin binding events lasting longer than 8 seconds. The number of PBAF binding hubs in each cell was reduced upon deletion of BAF180 bromodomains (Figure 2C). We saw a similar increase in the number of hubs detected for WT PBAF relative to ΔBD PBAF when we relaxed the stringency of our hub detection analysis by using PBAF chromatin binding events lasting longer than 1 second (Figure S4B). Correspondingly, the number of PBAF binding hubs increased in a BAF180 bromodomain-dependent manner when global levels of histone acetylation were elevated via SAHA treatment (Figure 2C and Supplemental Figure S4C).

Of note, removal of BAF180 bromodomains only weakened but did not completely eliminate ΔBD PBAF’s ability to bind chromatin hubs. This suggests that only a subset of target chromatin hubs are dependent on BAF180 bromodomains. Alternatively, additional chromatin binding domains (e.g. bromodomains, BAH, and PHD) in other PBAF subunits compensate for the removal of BAF180 bromodomains. Overall, our data indicate that at least a subset of target hubs are defined by repeated rounds of PBAF binding to chromatin via BAF180 bromodomains/acetyl-lysine interactions.

### BAF180 bromodomains do not inherently promote faster association of PBAF with chromatin hubs

Our data thus far indicated that bromodomain/acetyl-lysine interactions in nucleosomes strengthens PBAFs binding to chromatin, consistent with previous in vitro biochemical studies [12, 13, 25]. However, prior studies did not address if bromodomain/acetylated histone contacts increased the association (*k*_*on*_) or decreased the dissociation (*k*_*off*_) of PBAF binding to chromatin *in vivo*. We took multiple approaches to answer this question. To analyze PBAFs association with chromatin, we investigated PBAF cycling rates on chromatin in target hubs. Chromatin hubs are occupied via a series of PBAF binding and unbinding events interspersed with latent periods of non-occupancy (Figure 2D). If BAF180 bromodomain/acetylated histone interactions directly promoted faster association (e.g. increased *k*_*on*_) of PBAF with chromatin, histone hyperacetylation should increase WT PBAF’s cycling rates on chromatin. Histone hyperacetylation did not significantly impact cycling rates of WT or ΔBD PBAF on chromatin in hubs (Figure 2E). This indicates the histone hyperacetylation likely only increases the number of chromatin targets that are bound by PBAF’s BAF180 bromodomains while having little effect on promoting faster association with acetylated chromatin.

In addition, ΔBD PBAF should cycle less frequently on chromatin hubs compared to WT PBAF if BAF180 bromodomain/histone acetylation interactions promoted faster association (e.g. increased *k*_*on*_) with acetylated chromatin. In stark contrast with this idea, ΔBD PBAF exhibited more rounds of repeated binding events on hubs compared to WT PBAF when normalized to the total number of binding events in each cell (Figure 2E). Overall, differences in cycling rates between WT and ΔBD PBAF indicate that the time-dependent fluctuation of fluorescent signals in hubs is not due to inherent blinking of the dye which should be independent of the protein that is labeled. Rather, rapid cycling of PBAF on and off chromatin hubs is likely due to repeated rounds of binding and unbinding to chromatin.

The increased cycling rate of ΔBD PBAF on chromatin may be due to mass action effects since removal of BAF180 bromodomains allows ΔBD PBAF to target only a limited subset of chromatin hubs. In other terms, WT PBAF cycles less frequently on chromatin since it has more potential hub targets, essentially diluting out the concentration of PBAF available for re-visiting (see discussion). Consistent with this idea, the number of hubs targeted by ΔBD PBAF decreases 46%, while cycling frequency increases by 39% compared to WT PBAF. Thus, our data supports that BAF180 bromodomains do not promote faster association of PBAF with acetylated chromatin.

### PBAF stably engages chromatin in hubs via BAF180 bromodomains

*In vitro* biochemical experiments indicate that PBAF remodels/evicts nucleosomes via a multi-step pathway involving an initial transient encounter with a nucleosome (state 1), stable nucleosome engagement (state 2), and ATP-dependent nucleosome remodeling followed by dissociation of PBAF from the DNA scaffold (state 3) (Figure 3A). Based upon this mechanism, BAF180 bromodomains could promote efficient stable engagement of acetylated chromatin (e.g transition from state 1 to state 2). SMT is a powerful technique that measures kinetic parameters of different populations of chromatin bound factors [5, 20, 26]. Therefore, we analyzed PBAF’s chromatin binding residence times within hubs. Statistical analysis indicated that histograms of PBAF’s chromatin binding residence times inside of hubs were best fit with a double exponential decay model (Figure 3B). The predominant PBAF population (80% of molecules) transiently bound chromatin inside of hubs for 0.94 seconds (Figure 3B & Supplemental Figure S5). Transiently bound PBAF most likely represents non-specific or non-productive binding (Figure 3A, state 1) as seen previously with the Halo-tag alone and other transcription factors [20]. The remaining PBAF molecules (20%) stably bound chromatin inside hubs for 12.7 seconds further highlighting PBAF dynamic interaction with chromatin (Figure 3B). Importantly, PBAF’s stable residence time on chromatin (12.7 seconds) is significantly shorter than dye photobleaching rates (t_1/2_ of approximately 60-200 sec) indicating that these imaging conditions measure dissociation of PBAF from chromatin.

**Figure 3:**
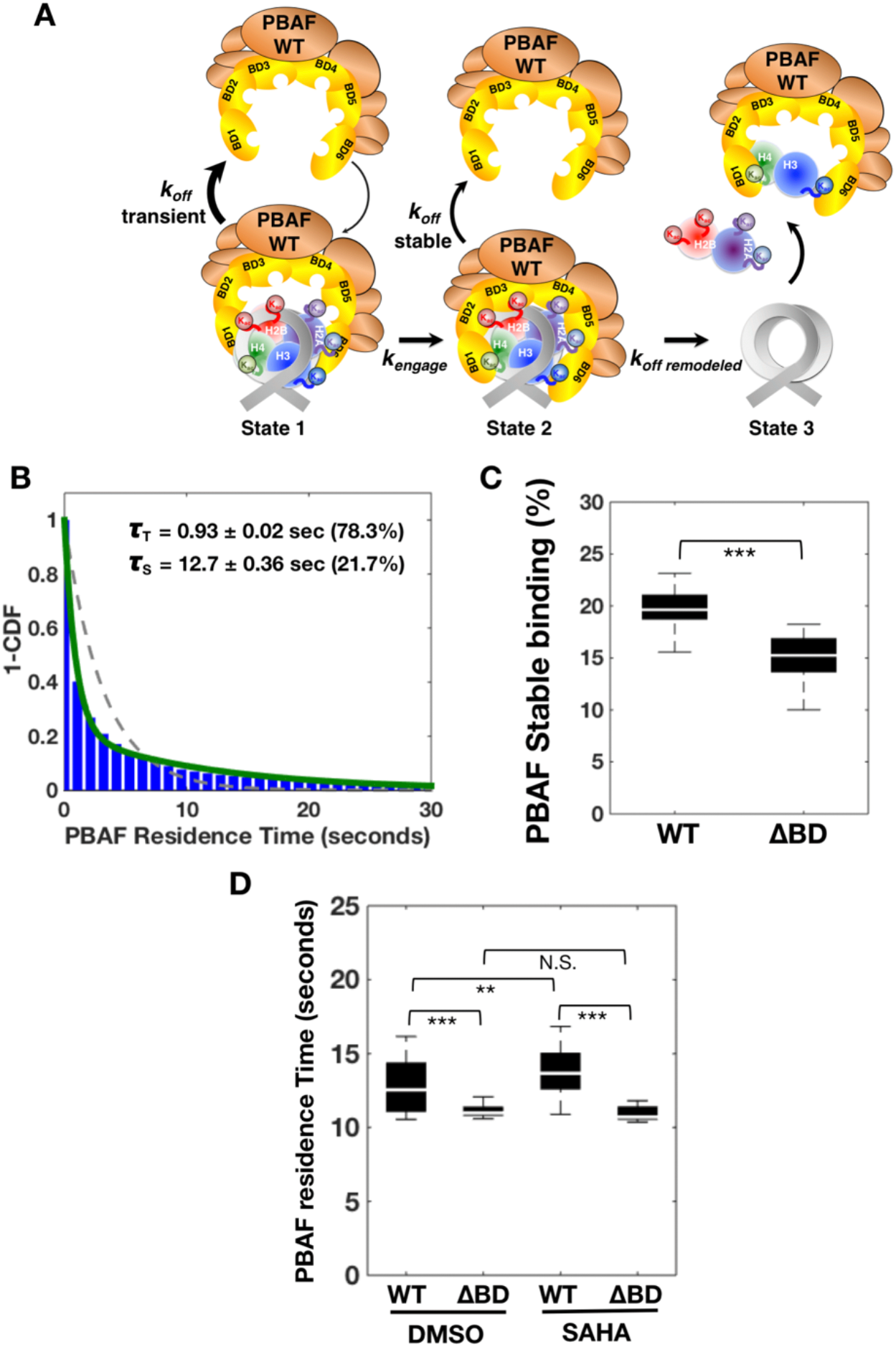
Spatial analysis of PBAF’s stability on chromatin *(A)* A schematic showing kinetic mechanism for PBAF binding and remodeling of a nucleosome. *(B)* 1−<u>C</u>umulative <u>D</u>istribution <u>F</u>unction Plot (1−CDF) of WT PBAF’s chromatin binding residence time inside hubs in a cell. 1-CDF plots were fitted to a single (dashed red) and double (solid green) exponential decay model. The residence time (τ) and its corresponding percentage for each population (i.e. Transient [T] and Stable [S]) are indicated. *(C)* Median percentage of Halo-BAF180 wild-type (WT) or Halo-BAF180 ΔBD (ΔBD) binding events displaying stable (S) binding to chromatin inside and outside of hubs (N=32 WT cells and N=31 ΔBD cells). *(D)* Median stable (S) chromatin binding residence time for Halo-BAF180 wild-type (WT) or mutant Halo-BAF180 ΔBD (ΔBD) inside of hubs (N=32 WT cells, 31 ΔBD cells). For data shown in *(C)* and *(D)*, the white bar in the solid black box is the median, while the lower and upper boundaries of the black box correspond to the 25th and 75th percentiles of the data respectively. (N=16 BAF180 WT/H3.3 cells, 16 BAF180 ΔBD/H3.3 cells, 16 BAF180 WT/ HP1α cells, and 15 BAF180 ΔBD/ HP1α cells). N.S. = not significant, p value ** = < 0.01 and *** = <0.001. Outliers typically represented less than 10% of the dataset and therefore were omitted for clarity.

To determine if BAF180 bromodomains promote stable association with acetylated chromatin, we compared WT and ΔBD PBAF’s chromatin binding residence times inside of hubs. Removal of BAF180’s bromodomains (ΔBD PBAF) decreased the percentage of PBAF molecules displaying stable binding inside hubs (Figure 3C). Therefore, BAF180 bromodomains enhance the efficiency of PBAF’s stable engagement with acetylated chromatin (Figure 3A, transition from state 1 to state 2).

Differences in residence times (**τ**_S_) within a population of molecules that are stably bound to chromatin directly reflect dissociation rates of PBAF from a nucleosome. To determine if BAF180 bromodomains decreased the dissociation rates (e.g. *k*_*off*_) of PBAF from chromatin, we compared the residence times (**τ**_S_) of WT and ΔBD PBAF inside hubs. Indeed, WT PBAF displayed longer stable residence times on chromatin compared to ΔBD PBAF (Figure 3D). Our data thus far indicates that once stably engaged on acetylated chromatin, BAF180 bromodomains decrease the dissociation rate of PBAF (Figure 3A, *k*_*off* *stable*_).

Hyperacetylation of histones (SAHA treatment) did not affect the percentage of stable chromatin binding events of WT or ΔBD PBAF inside hubs (compare Figure 3C and Supplemental Figure 6A). This suggests that histone hyperacetylation doesn’t further increase the efficiency of stable engagement of chromatin (Figure 3A, transition from state 1 to state 2). Histone hyperacetylation does slightly enhance the stability of PBAF once fully engaged on a nucleosome (Figure 3D). Therefore, histone hyperacetylation allows PBAF to find new target hubs in global chromatin (Figure 2C) and further stabilizes PBAF once bound to a nucleosome (Figure 3A, state 2).

### Chromatin inside and outside of hubs are distinct interaction scaffolds for PBAF

Chromatin hubs likely contain histone PTMs that better promote PBAF’s binding to nucleosomes relative to global chromatin. To test this hypothesis, we compared kinetic profiles of PBAF’s chromatin binding residence times inside and outside of hubs. Indeed, the percentage of PBAF molecules stably engaged with chromatin dropped precipitously outside compared to inside hubs. (compare Supplemental Figure S6A and Figure 3C). This suggests that chromatin in target hubs are likely decorated with distinct histone PTMs or exist in unique structures to promote the efficiency of PBAF’s engagement of a nucleosome compared to global chromatin outside of hubs.

Global chromatin outside of hubs could also present a unique scaffold that affects PBAF’s stability on chromatin. Therefore, we compared PBAF’s stable chromatin binding residence time outside and inside of hubs. PBAF’s stable chromatin binding residence time increases outside of hubs relative to inside of hubs (compare Supplemental Figure S6B and Figure 3D). Overall our data suggests that PBAF’s engagement is inefficient on global chromatin. However, once PBAF efficiently binds a nucleosome on global chromatin it is stabilized via unique histone PTMs and/or structures. This further suggests that chromatin inside and outside of hubs present unique scaffolds that regulate PBAF’s engagement and binding stability on a nucleosome.

Our data from inside chromatin hubs suggested that BAF180 bromodomains promoted efficient engagement and stable binding on acetylated chromatin (Figures 3C and 3D). To further determine how BAF180 bromodomains impacted PBAF’s stable interaction with global chromatin, we compared WT and ΔBD PBAF’s kinetic binding profiles on chromatin outside of hubs. Removal of BAF180 bromodomains further decreases ΔBD PBAF’s stable engagement and residence time on chromatin outside of hubs (Figures S6A and S6B). Overall our data suggests that global chromatin is also acetylated, albeit likely to some lesser degree compared to chromatin in hubs (see discussion).

### Prototypical markers for euchromatin (H3.3) and heterochromatin (HP1α) form dynamic hubs

Previous studies have established that PBAF binds nucleosomes at promoters of both transcriptionally active and repressed genes [14, 15, 27–29]. To better understand the different chromatin structures that PBAF targets in hubs, we dynamically mapped nuclear subregions associated with prototypical markers of euchromatin (H3.3-SNAP) and heterochromatin (SNAP-HP1α) (Supplemental Movies S2 and S3). Hubs containing dense high frequency H3.3-SNAP genomic interactions were found within distinct subregions in the nucleus (Figure 4A, left panel). Likewise, heterochromatic hubs harboring high frequency SNAP-HP1α/chromatin interactions were also observed with similar exchange kinetics as seen in previous FRAP studies (Figure 4A, right panel and Supplemental Figure S7) [30, 31]. Overall, this approach identifies distinct nuclear hubs where proteins associated with euchromatin and heterochromatin dynamically load and unload on the genome in a live cell.

**Figure 4:**
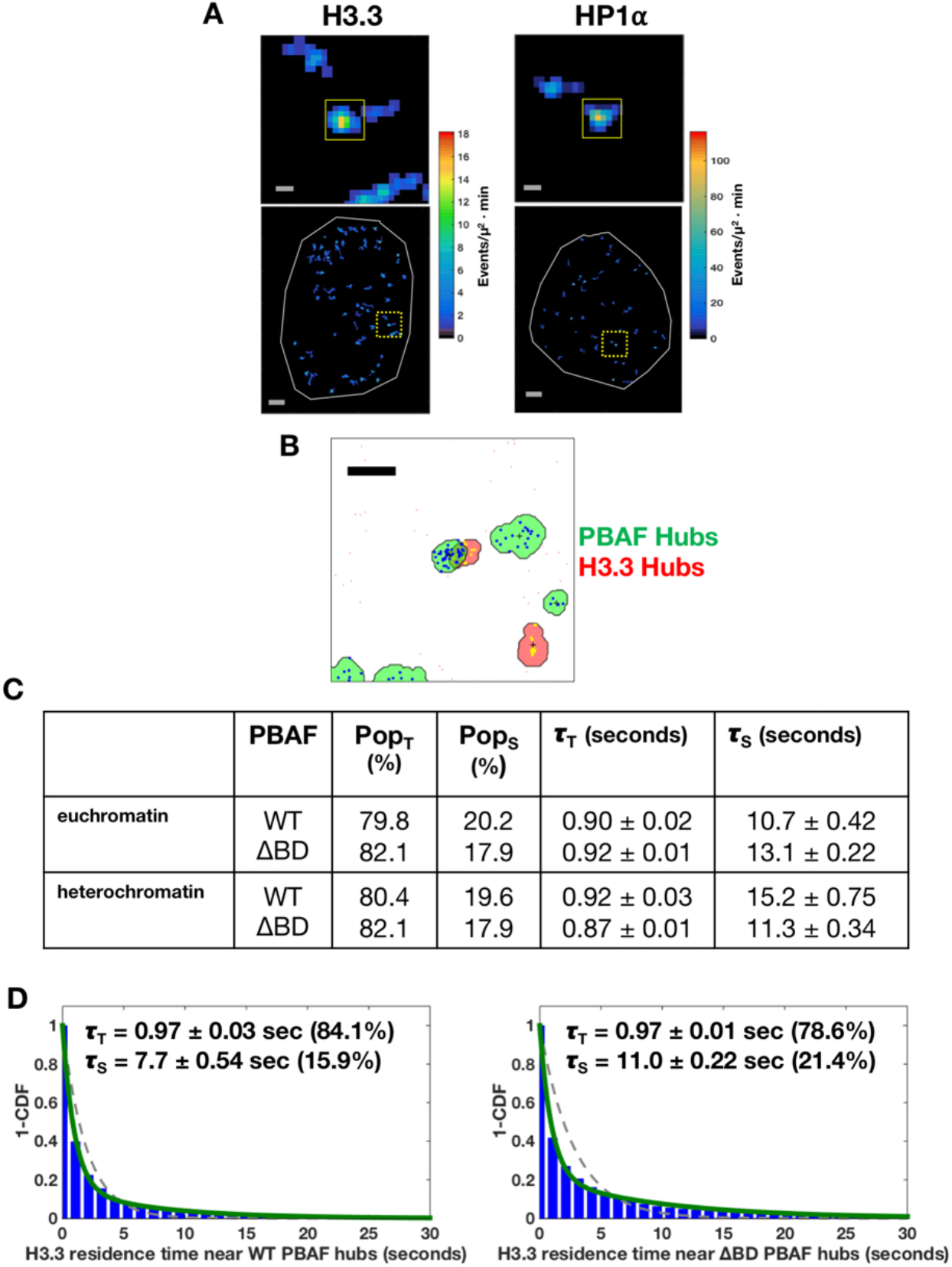
Spatial analysis of PBAF chromatin binding activities near H3.3 and HP1α marked chromatin hubs *(A)* High binding frequency hubs for H3.3-SNAP (left panels) or SNAP-HP1α (right panels). Top panels represent an enlarged view of the region outlined in yellow dashed boxes in bottom panels. The nucleus edge is shown in gray. Scale bars, 0.25 μm (top panel) and 2 μm (bottom panel). *(B)* Representative co-localization of a Halo-BAF180 WT hub (green hubs with blue binding events) and H3.3-SNAP hub (red hubs with yellow binding events). Scale bar, 0.5 μm. *(C)* Table of the distribution of transient (T) and stable (S) chromatin residence for WT and ΔBD PBAF near euchromatic and heterochromatic hubs. Median values for the population (Pop) are reported as percentages, while the residence time (**τ**) is in seconds. *(D)* 1−<u>C</u>umulative <u>D</u>istribution <u>F</u>unction Plot (1−CDF) of H3.3’s chromatin binding residence time near WT PBAF (left panel) or ΔBD PBAF (right panel) hubs in a cell. 1-CDF plots were fitted to a single (dashed red) and double (solid green) exponential decay model. The residence time (τ) and its corresponding percentage for each population (i.e. Transient [T] and Stable [S]) are indicated. N= 4,673 WT and 2,184 H3.3 binding events in 16 WT/H3.3 cells, 28,116 ΔBD and 15,130 binding events in 16 ΔBD/H3.3 cells, 2,563 WT binding events in 16 WT/ HP1α cells, and 8,859 ΔBD binding events in 15 ΔBD /HP1α cells. N.S. = not significant, p value * = < 0.05, ** = < 0.01 and *** = < 0.001. The white bar present in the solid black box is the median, while the lower and upper boundaries of the black box correspond to the 25th and 75th percentiles of the data, respectively. Outliers typically represented less than 10% of the dataset and therefore were omitted for clarity.

### PBAF’s stability on euchromatin and heterochromatin is differentially regulated via BAF180 bromodomains

To test if PBAF selectively engages different types of chromatin in hubs, we performed dual color single molecule imaging of PBAF (Halo-BAF180) and euchromatin ((H3.3-SNAP) or heterochromatin (SNAP-HP1α) in live cells. WT PBAF and euchromatic/heterochromatic hubs in close proximity (within 500nm) were identified and further analyzed to determine WT PBAF’s chromatin binding profiles (Figure 4B). WT PBAF did not selectively target euchromatic or heterochromatic hubs displaying no differences in percentages of WT PBAF molecules exhibiting transient versus stable binding of chromatin (Figure 4C). Therefore, PBAF’s stable engagement with chromatin hubs is independent of chromatin sub-type (e.g. euchromatic vs heterochromatic hubs).

When assaying PBAF’s stability on a bound nucleosome of different chromatin sub-types, we found that WT PBAF bound chromatin hubs near euchromatin (10.7 seconds) for significantly less time compared to heterochromatin (15.2 seconds) (Figure 4C).

Therefore, once bound to nucleosomes in hubs, PBAF’s chromatin bound residence time is dependent upon the type of bound chromatin structures. We speculate that PBAF’s, residence time may be dependent on whether the nucleosome needs to be evicted (transcriptional activation in euchromatin) or stabilized (transcriptional repression in heterochromatin) (Figure 5). This data is also consistent with WT PBAF’s enhanced stability on global chromatin which is predicted to be more stable and heterochromatic (compare 3D with Supplemental Figure 6B).

**Figure 5:**
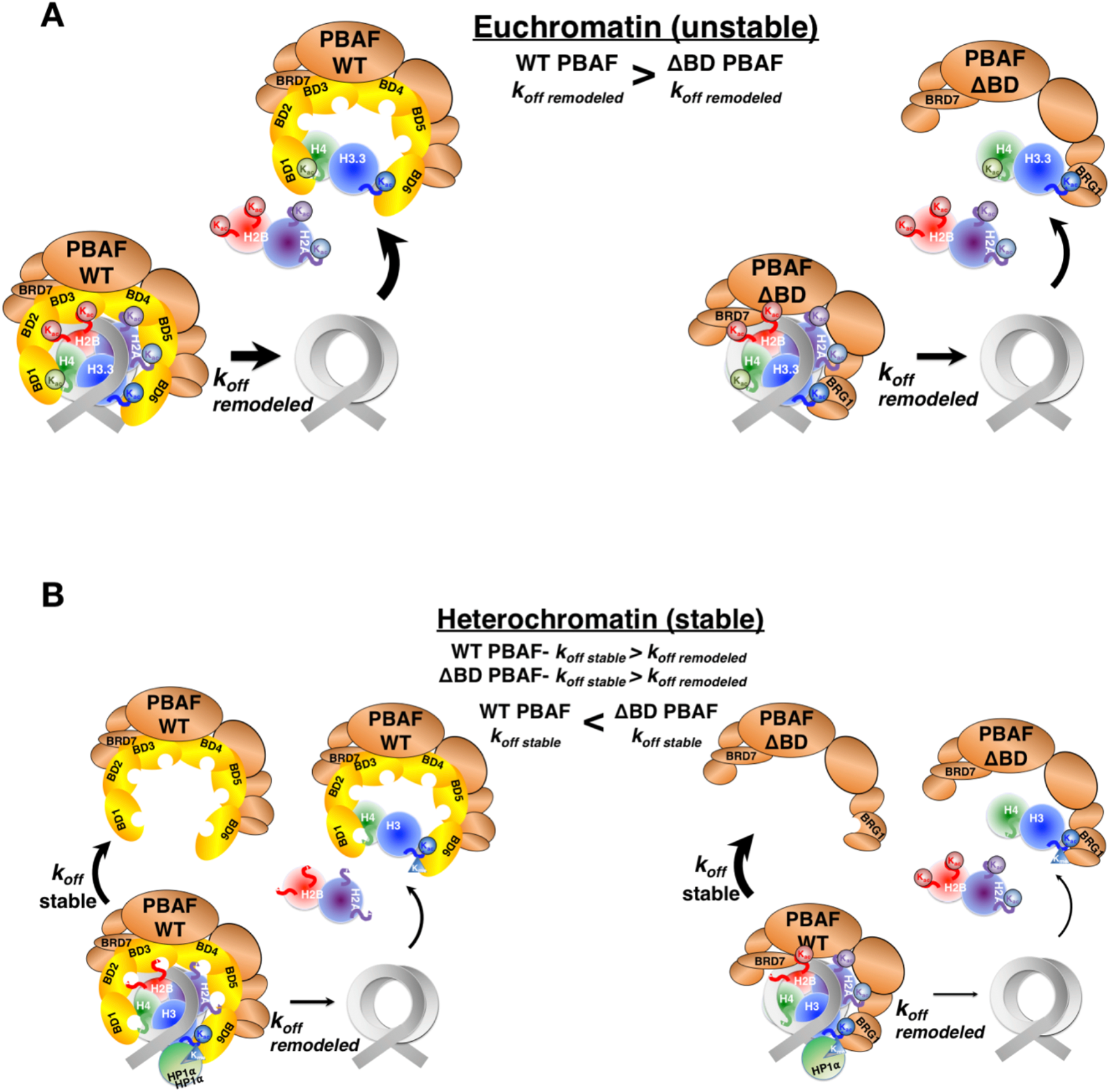
Models of PBAF’s dynamic chromatin binding to different chromatin states.*(A)* In euchromatin, PBAF’s main role is to evict or move nucleosomes to allow access of transcription factors to genomic sites. In this scenario, PBAF’s residence time is dominated by remodeling rates. Removal of PBAF’s bromodomains decreases the rate of remodeling (decreased *k*_*off* *remodeled*_) leading to a prolonged residence time in hubs. *(B)* By targeting heterochromatin, PBAF stabilizes a nucleosome for transcriptional repression or evicts a nucleosome to convert heterochromatin to euchromatin. In this scenario, PBAF’s residence time is can be reflected by either PBAF stabilizing (*k*_*off* *stable*_) or evicting (*k*_*off* *remodeled*_) a nucleosome. Since heterochromatin is likely remodeled slower than euchromatin, PBAF’s residence time may reflect dissociation from a non-remodeled nucleosome. As such, removal of PBAF’s bromodomains further reduces both the remodeling rates and stability on a nucleosomes leading to an increase in the rate of dissociation (*k*_*off* *stable*_).

Euchromatin is typically acetylation rich. Therefore, we predicted that BAF180 bromodomains would play an important role in stabilizing a bound PBAF complex to euchromatin. However, removal of BAF180 bromodomains led to an enhanced stability (e.g. decreased *k*_*off*_) of ΔBD PBAF bound near H3.3 hubs (e.g euchromatin) (Figures 4C). PBAF’s stability on chromatin is likely highly dependent on the ability of PBAF to remodel and evict the nucleosome (Figure 3A, see discussion). Therefore, it was possible that removal of BAF180 bromodomains effectively slowed down ΔBD PBAF’s ability to evict its bound H3.3 containing nucleosome resulting in an increased chromatin binding residence time. To test this hypothesis, we examined the residence times of H3.3-SNAP in close proximity (500nm) to either WT or ΔBD PBAF hubs. Consistent with our model, the stability of H3.3 on chromatin was higher near ΔBD PBAF compared to WT PBAF hubs (Figure 4D). Therefore, our data suggests that BAF180 bromodomains may help PBAF to rapidly disassemble H3.3 containing nucleosomes.

Residence time analysis of PBAF indicated that BAF180 bromodomains stabilized PBAF binding to global chromatin (Supplemental Figure S6B). This suggests that global chromatin contains acetylated chromatin. Since global chromatin is also likely highly heterochromatic, we compared the WT and ΔBD PBAF’s stability near HP1α hubs (e.g. heterochromatin). We find that ΔBD PBAF near heterochromatic hubs exhibited weakened stability on chromatin compared to WT PBAF (Figures 4C). This suggests that BAF180 bromodomains enhances PBAF’s stability on heterochromatin consistent with PBAF’s role in stabilizing nucleosomes to repress transcription [16, 17] (Figure 5B). Alternatively, BAF180 bromodomains stabilize PBAF on heterochromatin to help PBAF remodel a more stable nucleosome. To discern between these two possibilities, we measured the residence time of HP1α near WT and ΔBD PBAF hubs. Consistent with our model indicating that bromodomains help PBAF to remodel a nucleosome, the stability of HP1α on chromatin was higher near ΔBD PBAF compared to WT PBAF hubs (Supplemental Figure 7).

To support live-cell dynamic tracking of PBAF binding on select chromatin states, super-resolution STORM microscopy in fixed cells was also conducted (Supplemental Figure S8). The analysis validates the overlap between WT PBAF and H3.3/HP1α marked chromatin (Supplemental Figure S8, cyan). In many subnuclear regions, PBAF binds chromatin and forms a distinct close interface surrounding H3.3 marked chromatin domains (Supplemental Figure S8, bottom left panel, highlighted in red stars), consistent with the results in Figure 4B. A similar but less pronounced pattern was seen with PBAF and HP1α marked chromatin domains (Supplemental Figure S8, bottom right panel). Taken together, these studies illustrate that PBAF can selectively bind distinct types of chromatin via BAF180’s bromodomains.

## Discussion

A battery of single molecule imaging studies have now established that transcription is a stochastic process consisting of short bursts of transcription (minutes) intervened by periods of inactivity (minutes to hours). Transitions between transcriptionally active and inactive genes are likely accompanied by the surrounding chromatin interchanging between euchromatin and heterochromatin, respectively. Accordingly, this cyclical nature of going from active to inactive gene states suggests that chromatin remodeling enzymes, chromatin bound factors, histones and transcription factors dynamically bind and unbind chromatin at highly select regions of the nucleus over relatively short timescales. Indeed, our imaging data indicates that PBAF, H3.3 and HP1α can be added to the growing list of transcription factors (RNA Polymerase II, Mediator, EWS/FLI and Sox2) that have been shown to form hubs or clusters in live-cells [3–7]. Our imaging studies show discrete differences in the PBAF’s engagement and chromatin binding residence time inside and outside (Figures 3C, 3D and Supplemental Figures S5 and S6) of hubs further validating our hub calling approach. Therefore, we and other research groups have established that dynamic single molecule tracking is useful for identifying and characterizing chromatin binding of a broad range of nuclear factors in hubs of activity [7, 8].

However, little is known about how chromatin hubs are dynamically targeted and bound by chromatin remodelers. Using dynamic live-cell single molecule tracking, chromatin binding frequency heat maps allowed us to visualize PBAF binding to chromatin hubs. We propose that the discrete PBAF hubs identified in our heat maps (Figure 1D) represent repeated binding to arrays of closely spaced acetylated nucleosomes likely present in enhancers and promoters during our live-cell imaging (Figure 2). Spatial confinement of PBAF hubs over a small region of the nucleus (200-300nm) may also suggest that the genomic scaffold of chromatin targets does not move appreciable over our 18 minute imaging window.

Removal of BAF180’s bromodomains reduced the number of ΔBD PBAF hubs, while elevated global histone acetylation increased the number of hubs for WT PBAF (Figure 2C). Removal of BAF180 bromodomains did not completely eliminate ΔBD PBAF targeting to hubs (Figure 2C). Therefore, PBAF may utilize bromodomains in its BRD7/BRG1 subunits to target select BAF180 independent chromatin hubs. Alternatively, PBAF may also use additional chromatin binding domains (PHD and/or BAH domains) for engagement of BAF180 independent chromatin hubs.

Our cycling analysis reveals that removal of BAF180 bromodomains also increases ΔBD PBAF cycling on chromatin hubs relative to WT PBAF (Figure 2E). This increased cycling is likely due to a re-focusing of ΔBD PBAF engagement towards the remaining limited number of BAF180 independent chromatin hubs (Figure 2C). Finally, deletion of BAF180 bromodomains also weakened the efficiency of ΔBD PBAF’s engagement on chromatin by reducing the percentage of stably bound ΔBD PBAF (Figure 3C). This suggests that BAF180 bromodomains play a critical role in PBAF’s productive association (*k_engage_)* with chromatin (Figure 3A).

Different chromatin states can be, in part, defined by their nucleosome stability [32]. Unstable nucleosomes (e.g. acetylated euchromatin) are likely evicted from the genome by chromatin remodelers more easily and faster than stable nucleosomes present in heterochromatin. Thus, the residence time of chromatin remodeling enzymes on their nucleosomal targets may be highly dependent on the rate of chromatin remodeler induced histone eviction. Our imaging experiments support this idea by showing that PBAF displayed shorter chromatin binding residence times near H3.3 marked euchromatic hubs which should be more easily evicted compared to HP1α containing nucleosomes in heterochromatic hubs which should be less easily removed (Figure 4C). This is consistent with *in vitro* and *in vivo* studies showing an inherent instability of H3.3 containing nucleosomes [33–35]. Thus, rapid turnover of PBAF’s chromatin occupancy near euchromatin may directly reflect nucleosome stability which helps to define different chromatin states (Figure 5).

Based on in vitro experiments BAF180 bromodomains stabilize PBAF’s binding to acetylated chromatin. So one would naturally expect removal of BAF180 bromodomains would decrease ΔBD PBAF’s residence time on acetylated chromatin (e.g. euchromatin) in vivo. But these in vitro experiments were performed in the absence of ATP which prevents PBAF from evicting the nucleosome from DNA. So the authors were directly measuring BAF180 bromodomains affinity for acetylated chromatin. Our live-cell imaging studies are measuring PBAF’s chromatin binding and remodeling activity in the presence of ATP. As stated above, PBAF’s chromatin binding residence time in vivo is going to be a race between how long PBAF can hold on to acetylated histones (e.g. PBAF binding stability) and the time that it takes for PBAF to evict the nucleosome (e.g. nucleosomal stability) from the genome.

Contrary to the dogma of in vitro experiments, removal of BAF180 bromodomains increased ΔBD PBAF’s chromatin binding residence time near euchromatin. This suggests that once PBAF stably engages a nucleosome, BAF180 bromodomains don’t enhance the stability of PBAF on acetylated chromatin in vivo. This is consistent with a previous in vitro biochemical study of RSC, the yeast homolog of PBAF [25]. RSC, which contains multiple bromodomains, bound to an acetylated nucleosome with the same affinity as SWI/SNF, a highly related complex with only a single bromodomain. Therefore multiple studies now indicate that more bromodomains in PBAF/RSC doesn’t lead to stronger binding to acetylated chromatin.

Rather, ΔBD PBAF’s increased chromatin residence time may be a reflection of of a slower rate of nucleosome eviction. Indeed, our experiments showed that H3.3 is stabilized in chromatin when BAF180 bromodomains are removed (Figure 4D). Thus BAF180 bromodomains may help PBAF to speed up eviction of H3.3 containing nucleosomes in vivo (Figure 5A). Previous in vitro experiments also found that RSC stimulated nucleosome movement and H2A/H2B dimer displacement more than SWI/SNF in an acetylation dependent manner. Based upon our in vitro experiment and prior in vitro experiments, we speculate that removal of BAF180 bromodomains may be converting WT PBAF into a more “SWI/SNF like” complex with reduced chromatin remodeling activity.

Previous studies indicated that PBAF regulates transcriptional repression suggesting that PBAF also binds heterochromatin via an unknown mechanism [14–17]. Our live-cell and STORM imaging data (Figure 4B and Supplemental Figure S6) directly supports this premise that PBAF engages heterochromatin. Importantly, ΔBD PBAF’s chromatin binding residence time decreased near heterochromatin. This suggests that once PBAF engages heterochromatin, BAF180 bromodomains help to stabilize PBAF’s binding to a nucleosome. This BAF180 bromodomain mediated increase in stability on heterochromatin could be related to PBAF’s role in stabilizing a positioned nucleosome to repress transcription.

It is unknown how BAF180 bromodomains engage with heterochromatin that is typically envisioned to be deficient in acetylated histones. However, previous mass spectrometry studies have shown that chromatin contains bivalent modifications such as H3K9me3/H3K14ac on the same nucleosome [36, 37]s. We speculate that PBAF may recognize H3K9me3/H3K14ac in our HP1α marked heterochromatic regions given that HP1α binds H3K9me3 and the BAF180/BRG1 subunits interact with H3K14ac [19]. To increase PBAF’s stability on heterochromatin, it’s possible that PBAF simultaneously binds the H3K9me3/H3K14ac mark via the PHF10 PHD finger and the BAF180 bromodomains, respectively. Competition between PBAF and HP1α for H3K9me3/H3K14ac is supported by our data showing that removal of BAF180 bromodomains increased the stability of HP1α on chromatin (Supplemental Figure S7B). Additional future studies investigating the interaction of PHF10 and BAF180 with distinct chromatin marks inside cells should help test this hypothesis and further define PBAF’s engagement with heterochromatin.

Single molecule residence times have typically been interpreted to reflect binding affinities of transcription factors on target sites in chromatin. However for ATP-dependent chromatin remodeling enzymes like PBAF, a residence time is likely the time that it takes to evict/move a nucleosome, which is PBAF’s main role in vivo. In this regard, slowing down of PBAF’s enzymatic activity would slow down nucleosome eviction allowing PBAF to bind a nucleosome longer leading to an increased residence time.

This concept is consistent with our previous study showing that mutational inactivation of DNA Polymerase activity increased chromatin binding residence time *in vivo* [38]. This general phenomenon is likely due to the fact that an enzyme’s affinity for targets evolved to bind as long as it takes to exert its catalytic activity. However, when catalytic activity is inhibited, there is likely an upper time limit for substrate engagement leading to dissociation so that another enzyme can now attempt to utilize the substrate for catalysis.

## Conclusion

By characterizing the dynamic chromatin binding of PBAF and bromodomain mutants on different types of chromatin, our work helps to define the spatial and temporal changes of chromatin states. Future live-cell single molecule imaging studies of additional ATP-dependent chromatin remodelers along with histone PTM writers, readers, and the histone marks themselves will shine new light on the spatiotemporal organization of the 4D epigenome.

## Supporting information

Supplemental Movie S1 PBAF

## Acknowledgements

We thank Y.J. Chen and C.S. Peng for development of initial Matlab scripts used for SMT tracking. We are grateful to Z. Liu for providing the Matlab script and technical advice for analysis of diffusion rates and paired-correlation analysis. We thank S. Healton for reagents and technical assistance in acid extraction of histones, and J.C. Wheat for providing SAHA reagent and initial aliquots of acetyl-H3 antibody. We also thank D. Shechter for providing the histone H3 antibody. This work was supported by a grant from the NIH (8U01DA047729-04, RAC and RHS) as part of the 4D Nucleome project, the NIH (1R01GM126045-01, RAC) and a NIH Medical Scientist Training Program Grant (T32GM007288, CAK).

## Author contributions

Designed and supervised experiments: RAC, CAK, SHL, PC, WL, RHS

Generated material: CAK (cell lines, constructs), VW (cell lines, constructs), PD (cell lines), LDL (JF dyes)

Performed experiments: CAK, PC, VW

Analyzed data: CAK, SHL, PC, VW, RAC, WL

Wrote paper: RAC, WL, CAK

