## Supplemental Figures and Methods for "Bromodomains regulate dynamic targeting of the PBAF chromatin remodeling complex to chromatin hubs"

**Figure S1** **Kenworthy_Fig. S1**

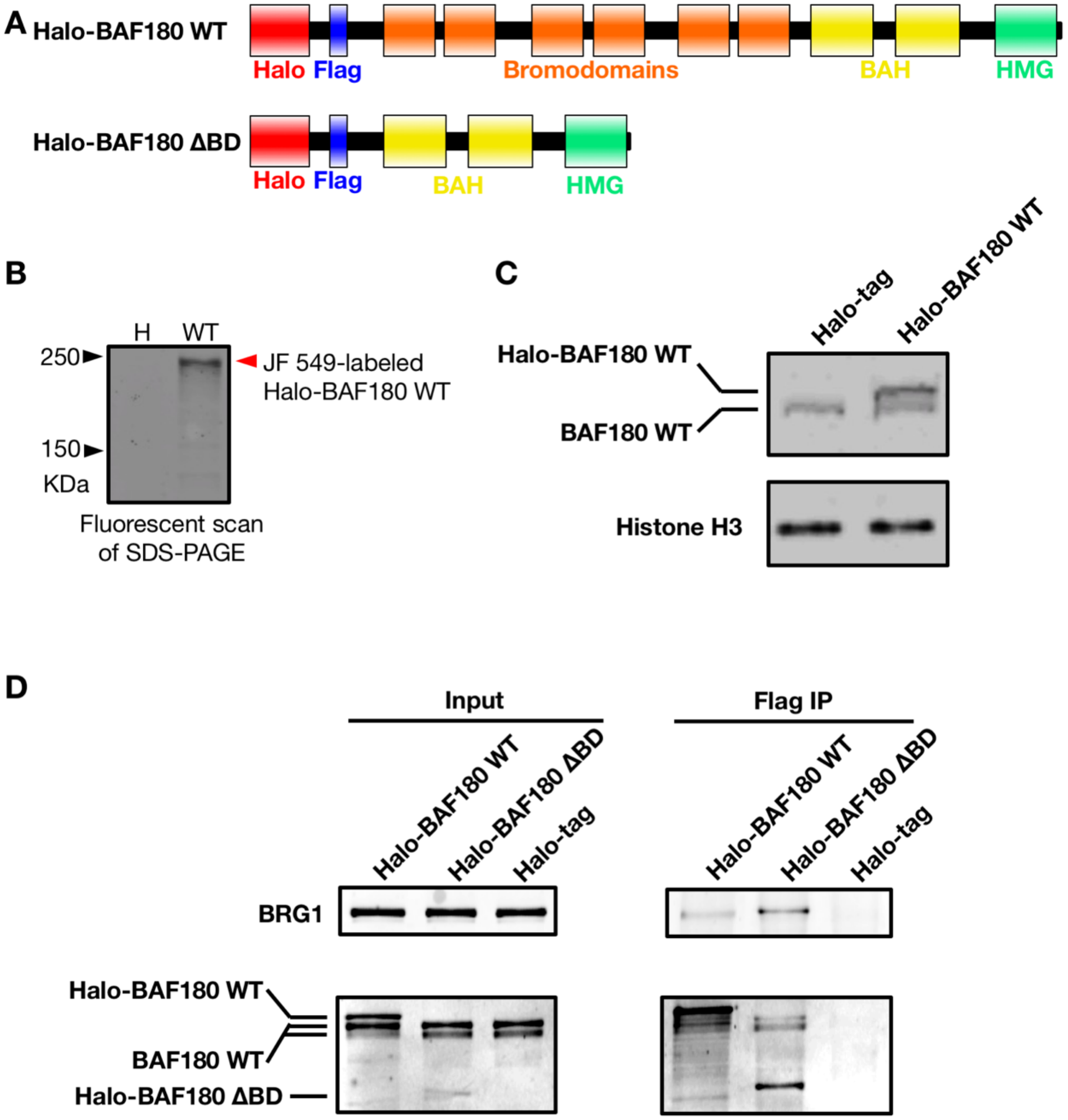


**Figure S1: Generation and characterization of stable U2OS cell lines expressing Halo-tagged BAF180, Related to Figure 1** *(A)* Domain diagram of wild-type Halo-BAF180 WT and mutant Halo-BAF180 ΔBD constructs generated for the current study. A Halo-tag and a flag-tag were incorporated into both constructs at the N-terminus of the protein. For Halo-BAF180 ΔBD, the N-terminal region before the first bromodomain was incorporated into the construct as this contained a bi-partite NLS to ensure transport of the protein into the nucleus. *(B)* Expression of the Halo-BAF180 WT construct was analyzed by *in vivo* labeling with JF549-HTL followed by SDS-PAGE. *(C)* Whole cell extracts generated from Halo-BAF180 WT and Halo-tag stable cells were analyzed by SDS-PAGE and western blotting using anti-BAF180 antibodies. A histone H3 western blot is provided as a loading control. *(D)* Nuclear extracts purified from Halo-BAF180 WT, Halo-BAF180 ΔBD and Halo-tag stable cells were immunoprecipitated using an anti-Flag epitope antibody, followed by western blotting using anti-HaloTag, BAF180 and BRG1 antibodies.

**Figure S2** **Kenworthy_Fig. S2**

**A**

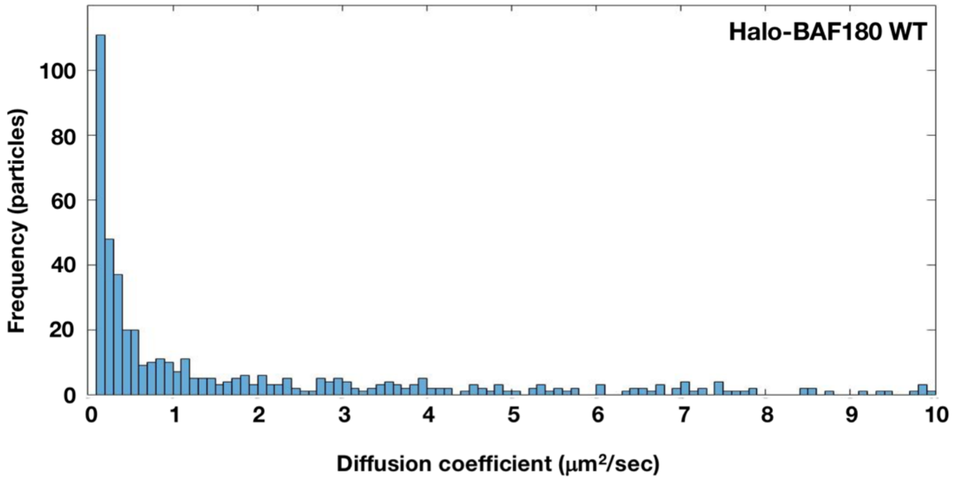


**B**

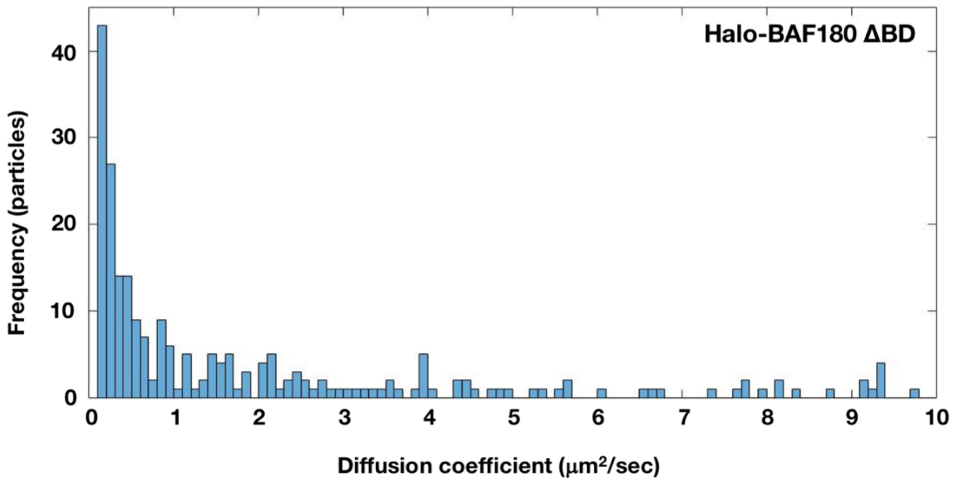


**Figure S2: Analysis of Halo-BAF180 WT and Halo-BAF180 ΔBD incorporation into PBAF *in vivo*, Related to Figure 1** *(A)* A probability distribution histogram of diffusion coefficients for freely diffusing Halo-BAF180 WT molecules labeled with JF646-HTL dye within the nucleus. The majority of PBAF molecules (71%) had a diffusion coefficient consistent with diffusion within a high molecular weight complex (1 MDa, 0.1-2 μm$^2$/sec). Importantly, only 10% of molecules had a diffusion coefficient consistent with freely diffusing Halo-BAF180 WT (180kDa, 3-5 μm$^2$/sec). This suggests that the majority of our measured localization events arose from Halo-BAF180 incorporated into a high molecular weight PBAF complex. *(B)* Free diffusion measurements were determined for U2OS cells expressing Halo-BAF180 ΔBD. Similar distributions were attained when compared to Halo-BAF180 WT expressing cells. 69% of molecules had a diffusion coefficient of 0.1-2 μm$^2$/sec consistent with its presence in a large molecular weight complex and 10% of molecules had a diffusion coefficient of 3-5 μm$^2$/sec.

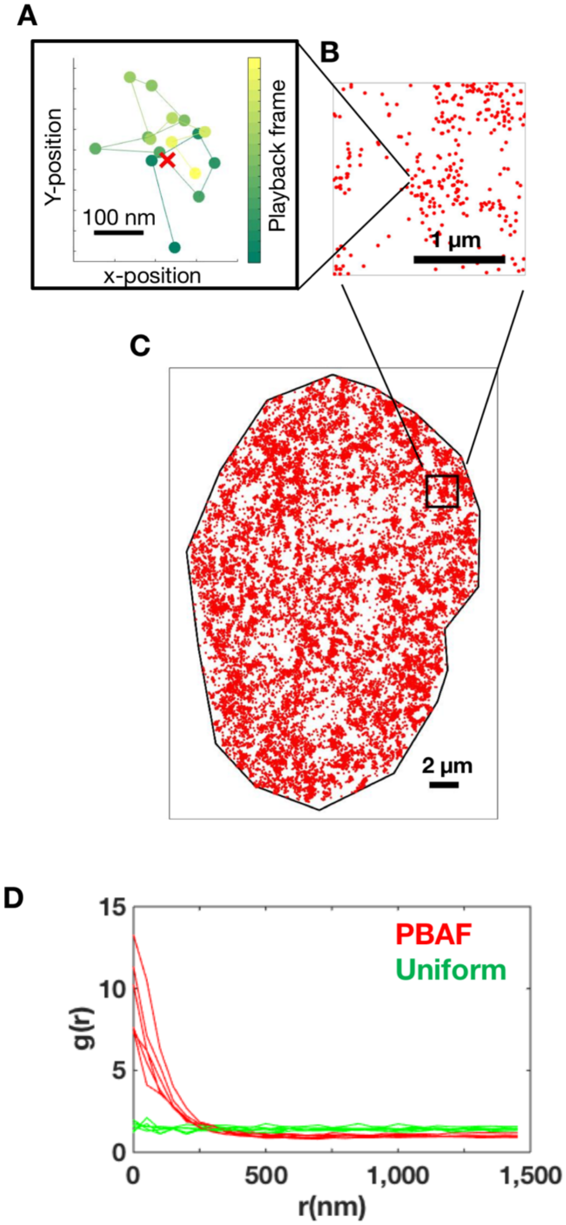


**Figure S3: Single molecule tracking and mapping of chromatin bound BAF180 in the nucleus, Related to Figure 1** *(A)* Localization of chromatin bound PBAF within a binding trajectory. X, Y localizations from 2D gaussian fits of PSFs (rainbow green dots) from consecutive frames of the movie were temporally linked based on a constrained diffusion model. Centroid location (indicated as a red colored X) of the binding trajectory was determined as the mean of x- and y-positions of the track. Scale bar, 100 nm. *(B)* Centroid positions (red dot) for each binding trajectory mapped within a nuclear subregion. Scale bar, 1 µm. *(C)* Centroid positions for each binding trajectory were mapped globally within the nucleus (outlined with black line). Scale bar, 2 µm. *(D)* Pairwise correlation analysis [1] of localization of PBAF binding events indicates clustering or hubs of PBAF binding at scales of around 200-400nm diameter. Bin size = 50nm, N = 6 WT PBAF cells

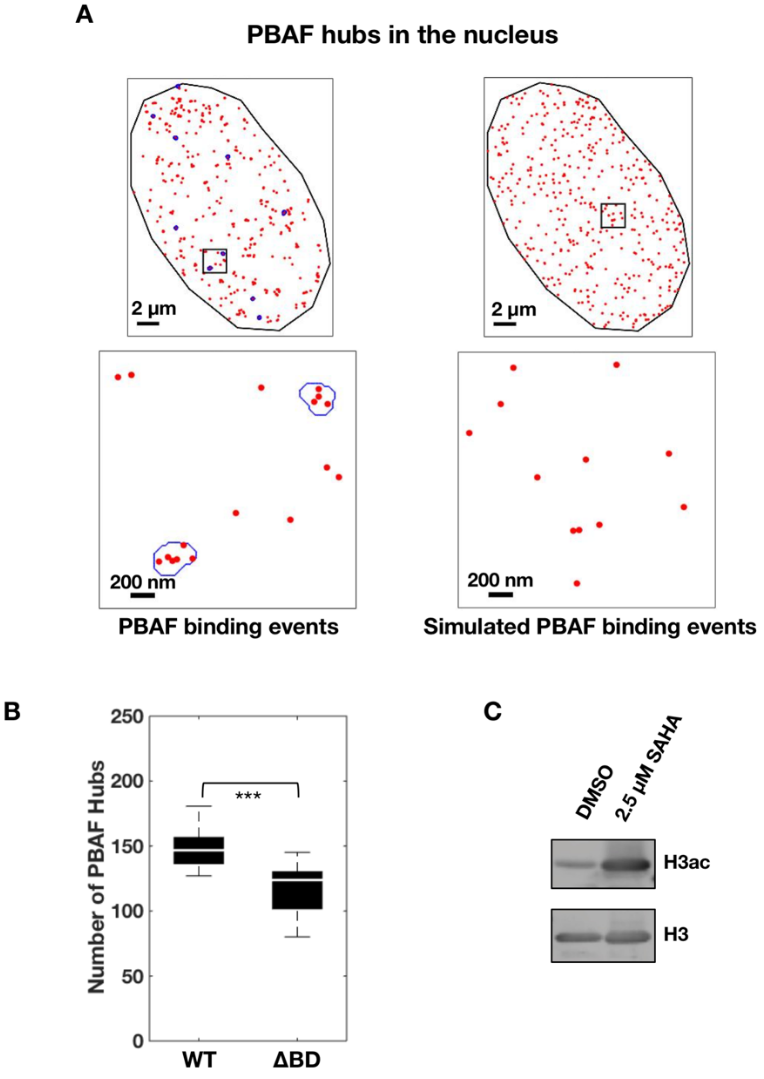


**Figure S4: Clustering analysis of genomic bound PBAF, Related to Figure 2** *(A)* Global wild-type (WT) PBAF chromatin binding maps were filtered based on duration of binding. PBAF chromatin binding events lasting longer than 12 seconds are shown in top left panel (red dots). Simulated PBAF chromatin binding maps shown in the top right panel (red dots) were generated by randomizing the positions of the filtered binding events (top left panel) within the nucleus. Clustering analysis algorithms revealed repeated PBAF binding events within small foci as hubs outlined in blue. Expanded insets (bottom panels) of boxed regions shown in top panels demonstrate clustering of PBAF binding events (red) within hubs (blue lines). Scale bars are indicated. *(B)* Clustering analysis using PBAF binding events lasting longer than 1 second. *(C)* Analysis of histone acetylation upon SAHA treatment of U2OS cells stably expressing Halo-BAF180 WT. Acid extractions from Halo-BAF180 WT expressing cells following 24 hours of 2.5 µM SAHA or equivalent vehicle treatment were analyzed by SDS-PAGE. Western blotting was performed against acetyl-H3 and normalized to total H3 as a loading control.

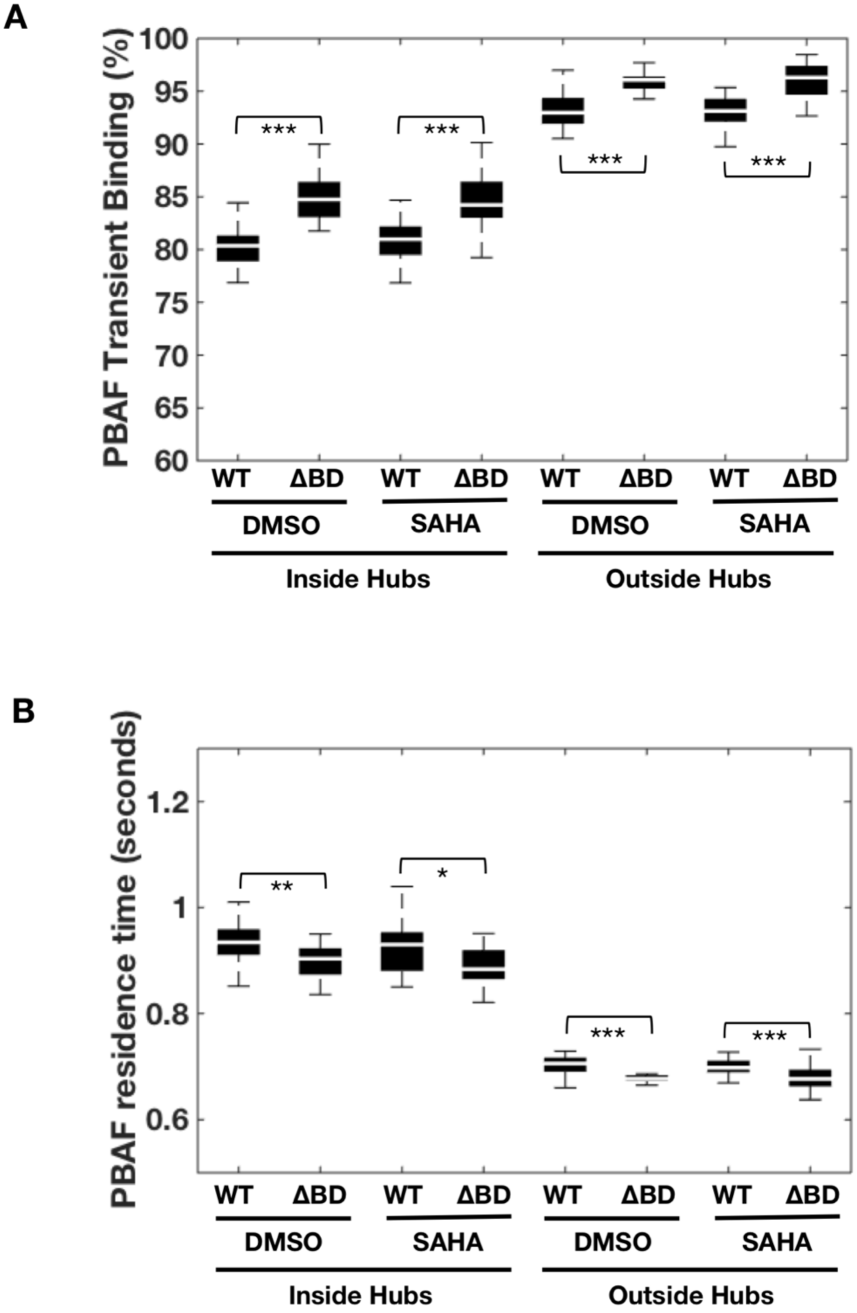


**Figure S5: Residence time analysis of PBAF transiently bound on chromatin inside hubs, Related to Figure 3** *(A)* Percentage of PBAF binding events inside hubs classified as the transient population via fitting of histograms of PBAF's residence time using a 2 component exponential decay model. *(B)* Residence time of PBAF molecules that are transiently bound to chromatin.

Figure S6 Kenworthy_Fig. S6

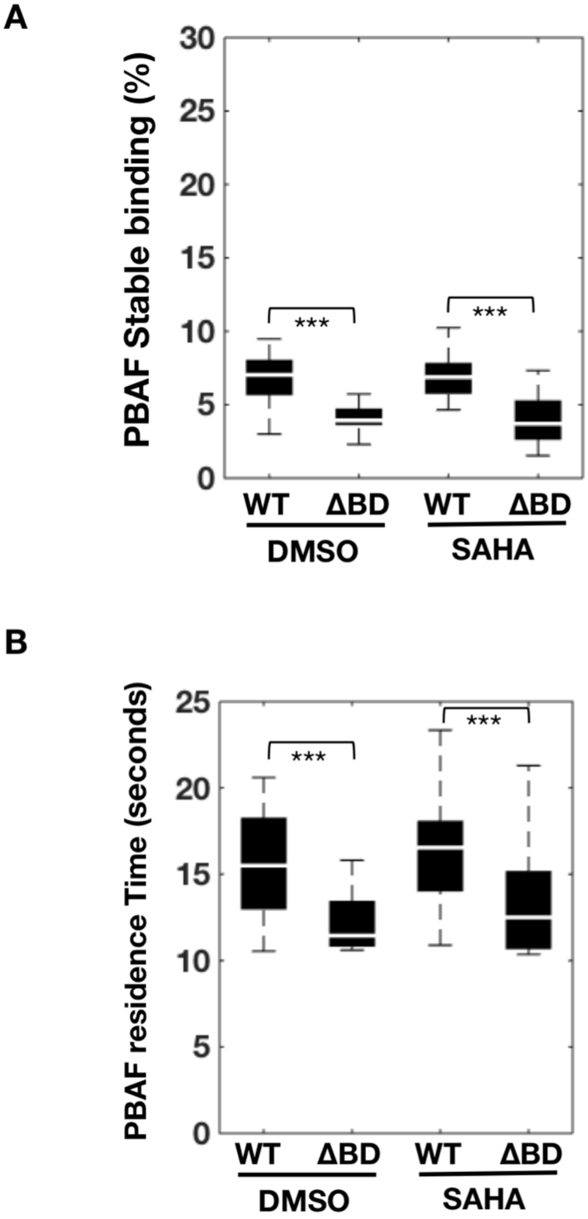


**Figure S6: Residence time analysis of PBAF stably bound to chromatin outside of hubs, Related to Figure 3** *(A)* Percentage of PBAF binding events outside hubs classified as the stable population via fitting of histograms of PBAF's residence time using a 2 component exponential decay model. *(B)* Residence time of PBAF molecules that are stably bound to chromatin.

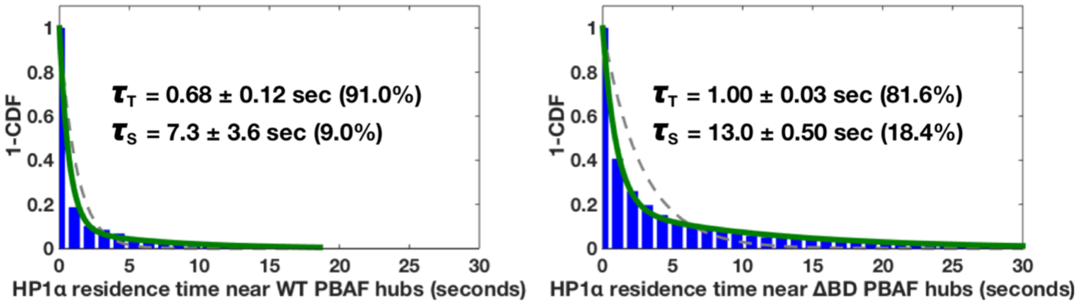


**Figure S7: Residence time analysis of HP1α bound to chromatin inside of hubs near PBAF hubs, Related to Figure 4** 1–Cumulative Distribution Function Plot (1–CDF) of HP1α's chromatin binding residence time near WT PBAF (left panel) or ΔBD PBAF (right panel) hubs in a cell. 1-CDF plots were fitted to a single (dashed red) and double (solid green) exponential decay model. The residence time (τ) and its corresponding percentage for each population (i.e. Transient [T] and Stable [S]) are indicated.

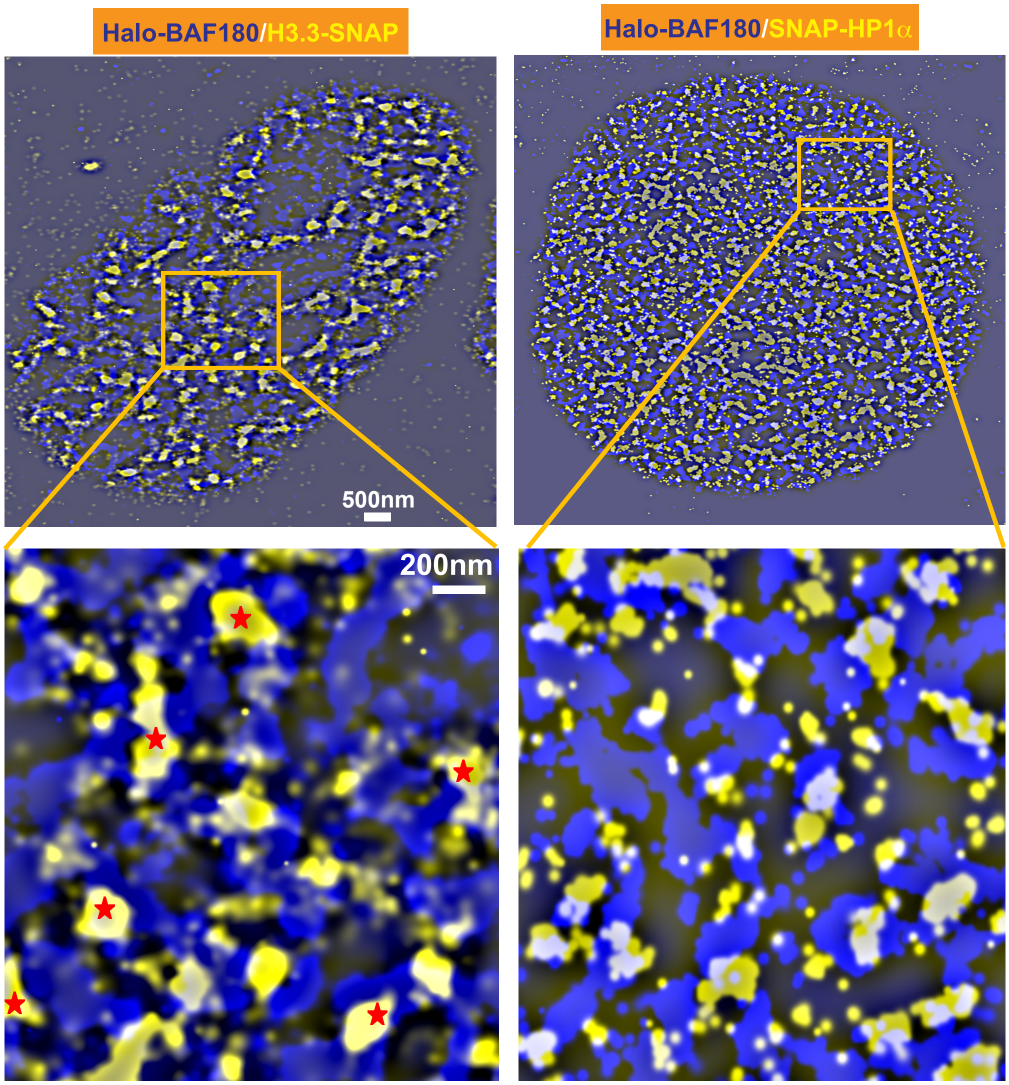


**Figure S8: STORM analysis of PBAF (Halo-BAF180), H3.3-SNAP, and SNAP-HP1α in U2OS cells, Related to Figure 4** Exposed nanodomains of Halo-BAF180 and Histone H3.3-SNAP or SNAP-HP1α (top panels). Super resolution imaging unveils discrete structures and interchromatin branching for BAF180 (blue). The protein intermixes with the two chromatin elements (histone H3.3 an HP1α shown in yellow) and exhibits cross sectioning with both target molecules (cyan). The bottom panels show filtered STORM images. Mild Fourier bandpass filtering exposes interface between BAF180 and the two chromatin interactors. Super resolution unveils a distinct preference of BAF180 to engulf H3.3 containing chromatin clusters in most cases from more than one side (asterisk marked H3.3 niches, bottom left panel). BAF180 exhibits a similar but slightly reduced proximity interface with HP1α containing chromatin clusters (bottom right panel). Scale bars are indicated.

### Supplemental Experimental Procedures

#### Cloning of Halo- and flag-tagged BAF180 construct

The wild-type human BAF180 coding sequence (isoform uc003deu.2) containing an N-terminal Flag tag and a C-terminal strep-tag II (i.e. BAF180 WT) was cloned into the pcDNA5-FRT-TOPO vector through PCR-amplification with the following complementary primers: forward primer 5'-ATGGACTACAAGGACGACGATGACA AAGGCCACCATCATCATCATCATTCTTCTGG-3' and reverse primer 5'-TATCCA CCTTTACTGTTACTTTTCAAATTGTGGGTGGCTCCAACTGCTCCCAACATTTTC TAGGTTGTATGCTTGGCG-3'.

A Halo-tag was added to the N-terminus of BAF180 WT by PCR amplification of the Halo-tag from the pENTR4-Halo plasmid (a gift from Eric Campeau, Addgene plasmid #29644) using the following primers: forward primer 5'-AGCGTTTAAACTTAAGCT CGCCCTTGCTCCACC-3' and reverse primer 5'-TGGCGGCGAGCTTAACAGTGGT TGGCTCGCCGG-3'. The Halo-tag PCR fragment was then inserted into the pcDNA5-FRT-BAF180 WT plasmid using the InFusion cloning system (Invitrogen). The Halo-tag alone was inserted into pcDNA5-FRT-TOPO through the TOPO cloning system.

#### Generation of the Halo-BAF180 ΔBD construct

Creation of the six bromodomain deletion construct (i.e. Halo-BAF180 ΔBD) was achieved on the pcDNA5-FRT-TOPO-Halo-BAF180 WT plasmid with mutagenesis primers (forward 5'-AGACGTAATCTCCAACATGGTACATAGTTGGAAGATTGGA AAGTCTCCTC-3' and reverse 5'-ATGTACCATGTTGGAGATTACGTCTATGTGGA ACCTGCAGAGGCCAACCT-3'). The resulting PCR containing amplified mutated circular DNA and template was then digested with Dpn1 overnight to remove original methylated plasmid template. The resulting template was transformed into DH5α *E. coli* competent cells. Clones lacking the six bromodomains were confirmed via restriction digest and sequencing analysis.

**Generation of the pSNAP-HP1α/H3.3 plasmids**

To generate pSNAP-HP1α plasmid, the HP1α coding sequence was cloned into pSNAPf plasmid (NEB) using geneblocks (IDT), so that the SNAP-tag was located at the N-terminus of the gene. To generate the H3.3-SNAP plasmid, the histone H3.3 coding sequence was PCR amplified from Pet28a-human H3.3 plasmid (a gift from Joe Landry, Addgene plasmid #42632) using the following primers: forward 5'-CGAGCTCGGATC GATATCGAATTGATGGCTCGTACA-3' and reverse 5'-TGCTGGCGCGCCGATAG CACGTTCTCC-3'. The resulting PCR fragment was then cloned into the pSNAPf plasmid through restriction digestion.

**Cell Culture and generation of FRT site, Halo-tag, Halo-BAF180 WT, Halo-BAF180 ΔBD, Halo-BAF180 WT/H3.3-SNAP, and Halo-BAF180 WT/SNAP-HP1α stable cell lines**

U2OS cells were grown in complete DMEM (high glucose DMEM supplemented with 10% FBS, 2 mM Glutamax [Fisher Scientific], 100 I.U./mL Penicillin, and 100 μg/mL Streptomycin [Corning]). To create U2OS cells containing a single FRT site (i.e. U2OS-FRT) for future generation of isogenic cell lines, we transiently transfected cells with pFRT/lacZeo plasmid containing an FRT site (Invitrogen). Single cell colonies were selected in 300 μg/mL zeocin. To generate cell lines expressing our Halo-tagged proteins, pFRT-Halo, pFRT-Halo-BAF180 WT, or pFRT-Halo-BAF180 ΔBD plasmids were co-transfected with pOG44 plasmid (Invitrogen) into U2OS-FRT cells. Selection for Halo-tag, Halo-BAF180 WT, or Halo-BAF180 ΔBD stably expressing cells was performed by supplementing media with 75-150 μg/mL Hygromycin B (Life technologies). To generate Halo-BAF180 WT/H3.3-SNAP or Halo-BAF180 WT/SNAP-HP1α stable cells lines, Halo-BAF180 WT stable cells were co-transfected with pSNAP-H3.3 or pSNAP-HP1α plasmids together with a second plasmid containing puromycin resistance. Cells co-expressing Halo-BAF180 WT along with either H3.3-SNAP or SNAP-HP1α were selected using 1μg/ml puromycin (ThermoFisher).

**JF549-labeling and SDS PAGE of U2OS-Halo and U2OS-Halo-BAF180 WT cells**

U2OS cells stably expressing the Halo-tag alone or Halo-BAF180 WT were incubated with 0.4 nM JF549 Halo-tag ligand (JF549-HTL) for 15 minutes at 37°C with 5% $CO_2$. Cells were then washed three times with 1x PBS before being incubated in complete DMEM for further 30 minutes at 37°C with 5% $CO_2$. Next, cells were harvested into 1x PBS containing 500 μM PMSF and pelleted at maximum speed for one minute before supernatant being collected and flash frozen in liquid nitrogen.

For SDS-PAGE, cell pellets were thawed on ice before being resuspended in 5x packed cell volumes (PCV) cytoplasmic extraction buffer (10 mM HEPES, 1.5 mM $MgCl_2$, 10 mM KCl, 500 μM PMSF, 10 μM Leupeptin, 1 mM DTT, pH 7.9) and incubated on ice for 15 minutes. NP-40 (0.2% final concentration) was added to lysates and tubes were inverted 5 times. Samples were centrifuged (2,000 x g) for 5 minutes at 4°C before cytoplasmic extract was transferred to a fresh tube. The remaining pellet was resuspended in 1x PCV nuclear extraction buffer (20 mM HEPES, 420 mM NaCl, 1.5 mM $MgCl_2$, 0.2 mM EDTA, 25% glycerol, 500 μM PMSF, 10 μM Leupeptin, 1 mM DTT, pH 7.9). Samples were then incubated on a nutator at 4°C for 15 minutes before being centrifuged (2,000 x g) for 20 minutes at 4°C. Supernatant was removed to fresh tube and supplemented with NP-40 to a final concentration of 0.1%. Soluble nuclear extract was quantified by Bradford assay. A total amount of ~7.4 mg protein for each nuclear extract sample was analyzed via a 7.8% SDS PAGE gel. Fluorescently labeled proteins (JF549) were visualized with a typhoon scanner using a 532 nm laser and a 650 nm long pass filter.

**Flag Immunoprecipitation and BRG1 western blotting**

The M2 flag resin (Sigma-Aldrich) was equilibrated in D100 wash buffer (10% glycerol, 2 mM $MgCl_2$, 20 mM HEPES, 100 mM KCl, 0.2 mM EDTA, 0.1% NP-40, pH 7.9). Cells stably expressing either Halo-tag alone, Halo-BAF180 WT, or Halo-BAF180 ΔBD were harvested prior to cytoplasmic and nuclear extractions as described above. BRG1 expression was assessed by Western blot analysis using a mouse monoclonal antibody against BRG1 (Santa Cruz; cat. No. sc-374197). Nuclear extracts were incubated with M2 flag resin for 2 hours on a nutator at 4°C. The resin was washed extensively with D300 wash buffer (10% glycerol, 2 mM $MgCl_2$, 20 mM HEPES, 300 mM KCl, 0.2 mM EDTA, 0.1% NP-40, pH 7.9) followed by D100 wash buffer. Drained resins were resuspended in 4x SDS loading buffer, boiled at 95°C for 2 min and run on a 7.8% SDS-PAGE gel. Western blotting was then performed on BRG1 followed by

BAF180 (Millipore rabbit polyclonal; cat no. ABE70) using fluorescently labeled secondary antibodies and visualized on a LI-COR Odyssey scanner.

**BAF180 western blotting of Halo-tag and Halo-BAF180 WT cell extracts**

Snap frozen pellets from cells stably expressing either the Halo-tag or Halo-BAF180 WT were thawed on ice before being re-suspended in 4x PCV NET buffer (150 mM NaCl, 5 mM EDTA, 50 mM Tris-HCl [pH = 8.0], 0.1% NP-40, 500 μM PMSF, 10 μM Leupeptin, 1 mM DTT) and were incubated on ice for 20 minutes. Lysates were then centrifuged (14,000 rpm) for 10 minutes at 4°C and supernatant was collected to a fresh tube as whole cell lysate. ~36 mg of lysate were loaded onto a 4-12% NuPAGE Bis-Tris gel (Invitrogen), and were run at 200V in 1X MOPS buffer. The gel was then transferred to a nitrocellulose membrane and western blotting was performed against BAF180 and Histone H3 (loading control) using fluorescently labeled secondary antibodies and visualized on a LI-COR Odyssey scanner.

**SAHA treatment, acid extraction, and histone Western blots**

Cells were seeded at a density of $1.6x10^6$ cells/$10cm^2$ dish overnight. After ~12hrs of growth, cells were treated with either 2.5 mM SAHA (Cayman Chemicals) or vehicle control (DMSO). After 24 hours, cells were harvested into 1X PBS plus 500 μM PMSF and were snap frozen in liquid nitrogen. Cytoplasmic and nuclear extractions were carried out as detailed above. Nuclear pellets were incubated in 100 mM $H_2SO_4$ overnight at 4°C. Extracts were then centrifuged (14,000 rpm) for 10 minutes at 4°C and supernatant was transferred to a new tube. Nucleosomes were precipitated by adding 2 mL of 100% ethanol for approximately 7 hours and centrifuging (14,000 rpm) for 10 minutes at 4°C. The pellet was re-suspended in NET buffer and protein content was quantified by Bradford assay. 10 mg of total acid extraction were run in duplicate on a 4-12% NuPAGE Bis-Tris gel (Invitrogen) in 1X MOPS buffer and transferred to a nitrocellulose membrane as detailed above. Western blotting was performed against Histone H3 and acetylated H3 (EMD Millipore, cat no. 06-599B) and visualized on a LI-COR Odyssey scanner.

**Live-cell fluorescent labeling of Halo-BAF180 WT and Halo-BAF180 ΔBD**

Cells stably expressing Halo-BAF180 WT were grown to a density of $\sim5x10^5$ cells on 35 mm MatTek imaging dishes. Twenty-four hours prior to transfection of Halo-BAF180 ΔBD, parental U2OS cells were grown to a density of $\sim1x10^5$ cells in 35 mm MatTek imaging dishes. Cells were then transiently transfected with the pFRT-Halo-BAF180 ΔBD plasmid and incubated overnight prior to further treatments.

Twenty-four hours before labeling, cells were treated with either 2.5 μM SAHA (Cayman Chemical) or matching vehicle control (DMSO, 0.25% final concentration) (Sigma) and were incubated at 37°C with 5% $CO_2$. Immediately prior to imaging, cells were incubated with 0.4 nM JF549-HTL dye for 15 minutes at 37°C. Cells were then washed 3 times with 1X PBS, replaced with complete DMEM and incubated for 30 minutes at 37°C to remove unincorporated dye. Cells were then washed 2 times with 1X PBS and placed in L-15 imaging media (Life technologies) plus 10% FBS for imaging.

**Dual color live-cell fluorescent labeling of Halo-BAF180 WT/ΔBD and H3.3-SNAP or SNAP-HP1α**

Cells stably co-expressing Halo-BAF180 WT along with either H3.3-SNAP or SNAP-HP1 were grown to a density of ~5x10$^5$ cells on 35 mm MatTek imaging dishes. Immediately prior to imaging, cells were incubated with 10 nM SNAP-Cell 647-SiR (New England Biolabs) and 0.4 nM JF549-HTL at 37°C for a total of 30 and 15 minutes, respectively. Cells were then washed 3 times with 1X PBS, replaced with complete DMEM and further incubated for 30 minutes at 37°C to remove unincorporated dye. Cells were then washed 2 times with 1X PBS and placed in L-15 imaging media (Life technologies) + 10% FBS for imaging. Two color labeling of Halo-BAF180 ΔBD along with H3.3-SNAP or SNAP-HP1α was performed as described above, except that Halo-BAF180 ΔBD were electroporated with pSNAP plasmids 24-72 hours prior to labeling.

**Halo-tag labeling and single molecule tracking (SMT) diffusion imaging**

Cells stably expressing Halo-BAF180 were seeded for ~15 hours, so that there would be ~5x10$^5$ cells upon imaging. Approximately 12 hours before imaging, cells were labeled for 30 minutes with 10 nM JF646-HTL dye before being replaced with complete DMEM and incubated overnight at 37$^o$C with 5% $CO_2$. The extended time between labeling and imaging was to reduce the background signal resulting from the free unreacted dye within the cell. Before imaging, cells were washed with 1x PBS and media was replaced with L-15 media (Gibco). Cells were then imaged at room temperature via a 640 nm laser (Coherent, ~103 W/cm$^2$), with a 256 x 256 pixel field of view and 2 x 2 binning, so that the effective pixel size was 168 nm. Cells were imaged for ~1.25 minutes with an effective acquisition time ≈ 25 ms.

**SMT diffusion analysis**

SMT analysis was conducted with MTT and custom MATLAB software as detailed in main methods. Tracks were then evaluated using the script evalSPT [2]. For analysis of diffusion, parameters were set as previously described [3] and tracks were examined using the DiffusionSingle matlab script (kindly provided by Zhe Liu). Mean Square Displacement (MSD) parameters were extracted for tracks from DiffusionSingle and those that were greater than 0.1 $\mu m^2$/second were evaluated.

**dSTORM Microscopy and analysis**

For dSTORM experiments, U2OS osteosarcoma cell lines stably expressing Halo tagged BAF180 and SNAP-tagged Histone H3.3 or SNAP-tagged HP1α were plated and cultured on 25 mm round coverslips (#1.5) for two days prior to imaging. The day of imaging, cells were labelled with Janelia Fluor 646 (JF646) (Tocris Biosciences) at 40 nM, along with SNAP-Cell TMR Star (NEB) at 100 nM, or with JF646 at 40 nM combined with SNAP substrate SNAP JF503 at 100 nM, in tissue culture medium (DMEM plus 10% FBS) for 30 minutes [4]. After this incubation period, cells were rinsed gently with plain medium 3 times, incubated with DMEM plus 10% FBS for 5 minutes in the tissue culture incubator and subsequently processed for fixation. Medium was aspirated and the cells were fixed in 1X PBS plus 4% paraformaldehyde (EM Grade Electron Microscopy Sciences at 15713) for 10 minutes at room temperature. Following fixation, the cells were rinsed twice with 1x PBS. For STORM imaging, we used on glass slides with single cavity well (Stellar Scientific), which was filled in with imaging buffer (50 mM Tris-HCl pH 8.0, 10% glucose, 50mM Cysteamine (Alfa Aesar #A14377), 0.5 mg/ml glucose oxidase (Sigma G7141), 40 μg/ml catalase (Sigma C1345) and the coverslips were mounted upside down. Excess liquid was removed making certain no air bubbles were trapped in the imaging chamber and the edges of the coverslip were sealed using epoxy resin. Imaging of each samples was done within the next 2-3 hours after chamber sealing. STORM imaging was performed on commercial NIKON super resolution systems, carrying an iXon Ultra EMCCD camera. Excitation was performed with 488 nm, 561 nm or 647 nm laser lines, depending on the combination of the fluorophores. Dual color labeling was performed in a simultaneous mode, but the recording was performed sequentially for the two fluorophores. The 647nm laser was used to record signal from the corresponding fluorophore, while the laser for the second fluorphore was set to zero. After acquiring 15,000-20,000 frames and almost completely bleaching the far red fluorophore, the 647 nm laser was tuned to zero power and the second excitation beam (488 nm or 561 nm) was used to collect data from the second fluorophore. This method minimized channel crosstalk and mechanical drifting. Data analysis was performed using the N-STORM (NIS-Elements) built-in analysis software. Drifting correction was done using cross correlation. In order to segment super resolved images and compare overlapping domains of BAF180 with Histone H3.3 or HP1α, images were filtered using ImageJ Fourier bandpass filtering large structures down to 40 pixels and small structures up to 3 pixels size. Pixel size on analyzed data was 2 nm. Filtered images were pseudocolored and overlayed using ImageJ plugins.
